# Fluorescence-based sorting of extracellular vesicles enables detection and omics

**DOI:** 10.64898/2026.10.07.753831

**Authors:** Marta Clos-Sansalvador, Marta Monguió-Tortajada, Sergio G Garcia, Eva Julià, Miriam Font-Morón, Erika Ramírez, Àlex Bote, Nicolás Baeza, Marta Sanroque-Muñoz, Linrong Pan, Berta Fusté, Julia Ponomarenko, Francesc E Borràs, Òscar Fornas, Marcella Franquesa

**Affiliations:** REMAR-IGTP Group Germans Trias i Pujol Research Institute (IGTP) & Nephrology Department, University Hospital Germans Trias i Pujol (HUGTiP), Can Ruti Campus Badalona (Barcelona), Catalonia, Spain; Department of Cell Biology, Physiology and Immunology Universitat Autònoma de Barcelona (UAB), Bellaterra, Spain; Universitat Pompeu Fabra, (UPF), Barcelona, Spain; Flow Cytometry Unit. Centre for Genomic Regulation (CRG), The Barcelona Institute of Science and Technology, Dr. Aiguader 88, Barcelona 08003, Spain; Hospital del Mar Research Institute, Barcelona, Spain; Department of Biochemistry and Molecular Biology, Universitat Autònoma de Barcelona (UAB), Bellaterra, Spain; Genomics Unit. Centre for Genomic Regulation (CRG), The Barcelona Institute of Science and Technology, Dr. Aiguader 88, Barcelona 08003, Spain; Bioinformatics Unit. Centre for Genomic Regulation (CRG), The Barcelona Institute of Science and Technology, Dr. Aiguader 88, Barcelona 08003, Spain; Department of Cell Biology, Physiology and Immunology Universitat de Barcelona (UB), Barcelona, Spain

**Keywords:** flow cytometry, FACS, exosome, extracellular vesicle

## Abstract

Extracellular vesicles (EVs) are nanosized, lipid bilayer-enclosed particles that mediate intercellular communication and hold considerable promise as biomarkers and therapeutic agents. However, their functional heterogeneity remains incompletely understood, largely due to the absence of reliable methods for isolating specific EV subpopulations.

Here, we established a Nanoscale Flow sorting (NanoFACS) workflow and evaluated its potential for EV detection, sorting, and molecular characterization. We assessed critical parameters, including fluorescent staining and labeling approaches, background signal discrimination, dilution and flow-rate effects, and scatter triggering.

We achieved high-purity sorting of CFSE-stained and antibody-labeled EVs, preserving vesicle integrity and sufficient RNA yields for small RNA sequencing of sorted populations. We identified EV subtype-specific small RNAs despite partial cargo overlap in single-marker–sorted populations.

Our results establish a reproducible antibody-based NanoFACS workflow that defines key operational parameters for reliable EV detection and sorting, with potential adaptability across conventional and spectral flow cytometry platforms. By reducing background and non-EV-associated events, the workflow enables the isolation of phenotypically defined EV subpopulations suitable for downstream molecular analysis, providing a practical approach to investigate EV heterogeneity.

**Graphical abstract:** 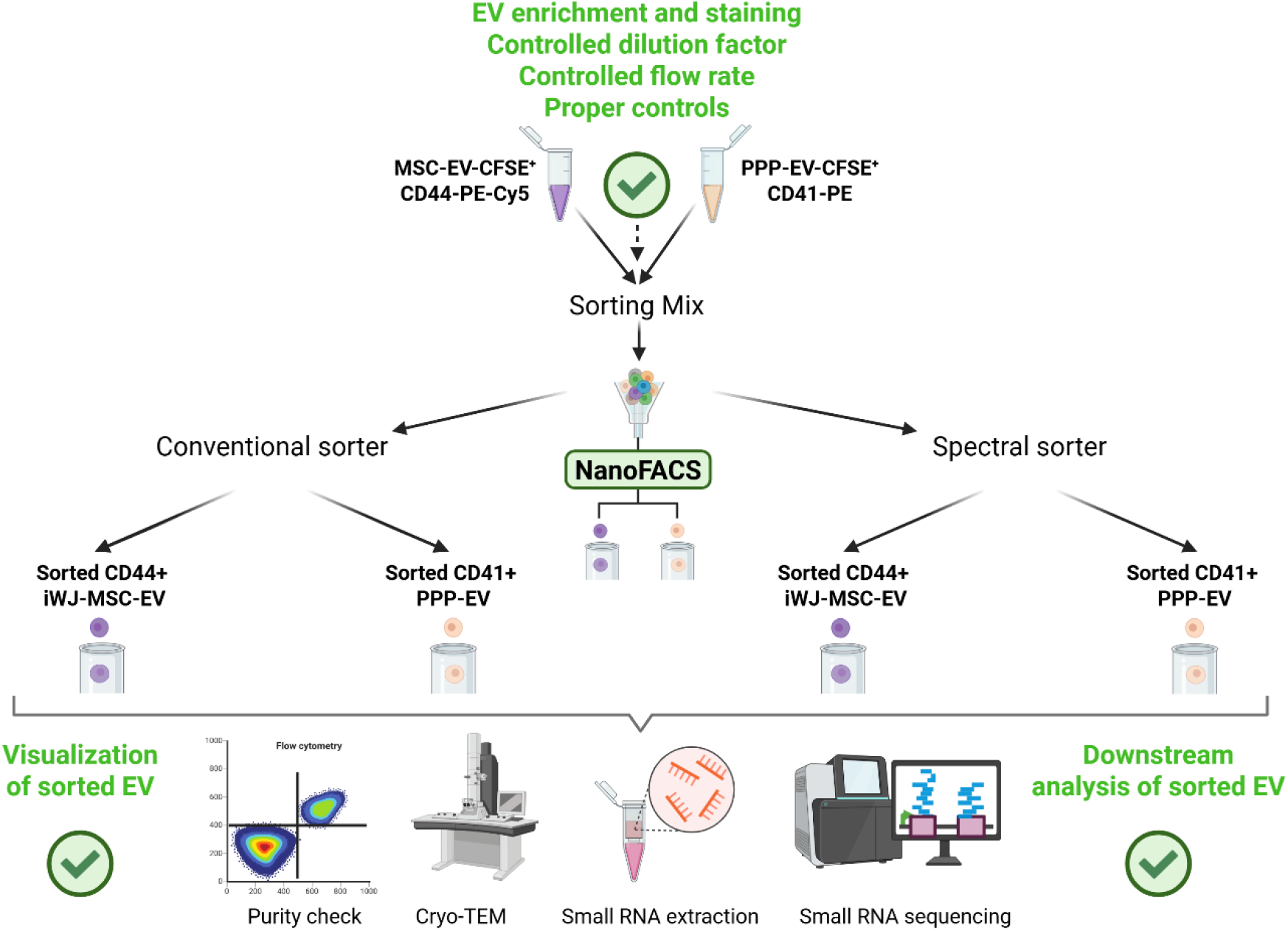

## Introduction

Extracellular vesicles (EVs) are complex, nanoscale biomaterials composed of a lipid bilayer enriched with membrane proteins and glycans encapsulating an aqueous core that contains protected nucleic acids, metabolites, proteins such as cytokines or other soluble mediators with functional bioactivity (1). Their intrinsic role in intercellular communication, along with their unique molecular cargo and biocompatibility, renders them highly attractive as source of biomarkers and for use as therapeutic agents, being a powerful biological tool in both diagnostics (2–4) and therapeutic applications (2,5).

EV heterogeneity encompasses variability in size, surface markers, cargo composition, and biological activity, even among EV secreted by a single cell type under uniform conditions (6). This diversity implies that the biological effects observed in EV preparations likely reflect the collective activity of distinct EV subtypes, rather than a homogeneous population. As such, specific functional outcomes may be mediated by rare or specialized vesicle subpopulations, which are diluted and potentially masked in pooled EV isolates. Likewise, EV populations from specific cells of origin, bearing useful biomarkers for diagnosis and/or prognosis can be scarce in complex biofluids like urine or plasma. To fully harness the potential of EVs as biomarkers and elucidate their roles in physiological and pathological processes, it is essential to sort them into well-defined subpopulations based on their cellular origin and molecular phenotype, with carefully characterized yield and purity. Therefore, there is an urgent need for high-resolution tools capable of isolating, profiling, and interrogating defined EV subtypes.

In this context, flow cytometry offers a promising avenue for addressing this challenge. Originally developed for single-cell analysis, fluorescence activated cell sorting (FACS) has been adapted for nanoscale particle sorting with increasing precision, proving especially useful for the distinction of viral populations (7) and nanoplastics (8). Until recently, only two studies have reported attempts to sort EVs by flow cytometry (9,10), yet with relative success as the sorted sample resulted highly diluted and was unsuitable for downstream applications or molecular analysis. Morales-Kastresana et al. (9) demonstrated the capability of nanoparticle sorting using a the MoFlo Astrios EQ system (Beckman Coulter) while Kormelink et al. (10) used the BD Influx (Becton Dickinson), both systems requiring highly skilled technicians and complex alignment modifications along with specific set-up protocols. Furthermore, these two studies did not seek to sort based on individual antibody-based labeling. Earlier this year, Ragni et al. (11) used a modified MoFlo Astrios to sort EV based on fluorescent staining only and performed antibody-based phenotyping and RNA extraction in the post-sorting fractions. Advancing EV-adapted FACS workflows is thus pivotal for unraveling EV functional heterogeneity. By leveraging fluorescent labeling of EV surface markers or cargo, flow cytometry offers a promising tool for high-throughput, quantitative sorting of individual vesicles based on molecular phenotype. This strategy would allow for the enrichment of discrete EV subpopulations, facilitating downstream analysis of their composition, function, and therapeutic potential. Nevertheless, technical hurdles remain, including the need for improved sensitivity, standardized labeling protocols, and robust controls to distinguish true EV signals from background noise or artifacts. In this line, very recently Castrosín et. al launched a preprint (12) assessing EV sorting using the Cytoflex SRT (Beckman Coulter) configuration to successfully sort EV by targeting and phenotyping them through violet laser (405nm) thresholding and using protein-specific fluorescent tags.

In the present manuscript we aimed at setting up a reliable method of nanoparticle FACS (nanoFACS) to consistently isolate single nanovesicles based on CFSE staining and antibody-based labelling of subpopulation-specific EV surface markers. We first used a customer modified conventional BD Influx to replicate the published evidence (10) and optimize a nanoFACS workflow as a proof-of-concept, which was further advanced to a user-friendly BD FACSDiscover™ S8 spectral imaging cell sorter (BD Biosciences). Specifically, we were able to sort CFSE-stained and antibody-labelled EV populations based on fluorescent antibody signal in both platforms. We followed MiFlowCyt-EV standards (13) of analysis and reporting, and used calibration material, proper EV controls and specific software to test the resolution capacity of the two cytometers and the feasibility of our approach based on fluorescent thresholding. The presence and integrity of sorted EVs were confirmed by electron microscopy and the cytometer imaging module. Finally, we performed smallRNA extraction and sequencing of the sorted EV populations, serving as a proof-of-concept of our nanoFACS workflow for the obtention of homogeneous EV populations in high purity and yield to allow molecular downstream applications.

## Materials and Methods

### Cell culture

Immortalized Wharton’s Jelly-derived Mesenchymal Stromal Cells (iWJ-MSC) were cultured in T-175 flasks using alpha-minimum essential medium (alpha-MEM; Sigma Aldrich, St Louis, MO, USA), supplemented with 10% heat-inactivated fetal bovine serum (FBS; Life Technologies, Carlsbad, CA), 0.02 mM of L-glutamine and 1% penicillin– streptomycin (P/S; Gibco Invitrogen Corp). The cultures were incubated at 37°C in a humidified atmosphere with 5% CO2. Non-conditioned cell culture media was used as a control and underwent the same sample processing as the MSC-conditioned cell medium. 100 kDa concentrated medium aliquots were frozen until extracellular vesicles (EV) isolation,

### Platelet-poor plasma (PPP) obtention

To obtain the platelet-poor plasma (PPP), 9 mL of blood from two different female healthy volunteers were collected under fasting conditions, previous informed consent, in sodium citrate anticoagulation tubes and gently mixed by inverting the tubes 3 times. Next, the sample tubes were centrifuged at 350 ×g for 20 min to pellet erythrocytes and absence of hemolysis was confirmed visually. Finally, the top layer of supernatant was collected and centrifuged at 2000 ×g for 10 min at room temperature to obtain the PPP. Supernatant was aliquoted and frozen at -80°C in 1 mL aliquots until EV isolation.

### Extracellular vesicles isolation

iWJ-MSC-EVs and PPP-EVs were isolated by size exclusion chromatography (SEC) following previous published protocols (14). Supernatant from 25×10^6^ - 35×10^6^ producing cells was cleared from cells and debris by centrifuging for 5 min at 400 ×g, and 10 min at 2000 ×g, respectively. Then, 45 mL of cleared supernatant or 2 mL PPP were concentrated using a 100 kDa molecular weight cut-off (MWCO) ultrafiltration unit (Amicon Ultra filter, Millipore, Billerica, MA, USA). 100 µL of concentrated iWJ-MSC supernatant or PPP were then fractionated by SEC in 1 mL-columns packed with Sepharose CL-2B (Sigma–Aldrich, St. Louis, USA) in 100 µL-fractions. Phosphate buffer saline (PBS; REF: 12899712; Fisher BioReagents) was used to elute the fractions, and EV contained in fractions 6 to 8 were pooled after bead-based assay confirmation. Pooled fractions were stored for a maximum of 24 h at 4°C prior to sorting experiments.

### Bead-assisted flow cytometry

The presence of EVs in SEC fractions was assessed using bead-based flow cytometry (15). Briefly, 10 μL of each SEC fraction were incubated with 2 μL of 4 μm-aldehyde/sulfate latex beads (Invitrogen-Thermo Fisher Scientific) for 15 minutes at room temperature (RT). The reaction was then blocked by adding 1 mL of BCB buffer (PBS containing 0.1% bovine serum albumin (BSA) and 0.01% sodium azide (NaN3); Sigma Aldrich) and incubated for 2 hours on a rotating platform at 15 rpm at RT. Subsequently, EV-coated beads were centrifuged at 2000 ×g for 10 minutes and resuspended in BCB buffer.

Antibody labeling was carried out by incubating the beads for 30 minutes at RT with either the fluorochrome-conjugated antibodies anti-CD90-PE-Cy7 (1/50; REF: 328124; BioLegend, Inc., San Diego, CA, USA) and CD41-PE (1/50; REF: 303706; BioLegend, Inc., San Diego, CA, USA) or by indirect labeling. For indirect labeling, primary antibodies anti-CD63 (1:10, Clone TEA3/18) and anti-CD9 (1:10, Clone VJ1/20.3.1) kindly gifted by Dr. Yáñez-Mó (Departamento de Biología Molecular, UAM/IIS-IP) were used, followed by secondary antibody incubation with FITC-conjugated goat F(ab’)2 anti-mouse IgG (1:100; Bionova, Halifax, NS, Canada) or Alexa647-conjugated donkey F(ab’)2 anti-mouse IgG (1:100; Jackson ImmunoResearch Europe Ltd., Ely, UK) in CFSE-labeled EVs. After each incubation step, the beads were washed with BCB buffer by centrifugation at 2000× g for 10 minutes.

Finally, data were acquired using a FACSLyric flow cytometer (Becton Dickinson, San José, CA) and analyzed with FlowJo v10.2 software (FlowJo, LLC, Ashland, OR, USA).

### EV staining

Fluorescently labeled iWJ-MSC-derived EVs were prepared for sorting. In brief, 100 μL of concentrated iWJ-MSC supernatant was incubated with 20 µM carboxyfluorescein succinimidyl ester (CFSE; eBioscience™, Invitrogen) for 2 hours at 37°C prior to EV isolation. Excess CFSE dye was removed through the subsequent SEC purification of the labeled EVs.

As a control of unspecific CFSE signal, non-supplemented alpha-MEM medium (devoid of EVs) was concentrated, stained with CFSE, and eluted through a SEC column as performed with iWJ-MSC-EV-rich supernatant.

### EV labeling

Typically, 200µL of undiluted MSC-EVs and PPP-EVs were incubated for 30 min with either the CD44-PE-Cy5 antibody (1/50; REF: 103010; BioLegend, Inc., San Diego, CA, USA) or the CD41-PE (1/50; REF: 303706; BioLegend, Inc., San Diego, CA, USA) antibody, respectively. Next, 1 mL of filtered (0.2 µm) PBS was added to each mix, loaded into an Amicon Ultra filter (Millipore, Billerica, MA, USA) of 10 kDa and centrifuged for 15 min at 3000 ×g to wash the unbound antibody. Finally, the washed and labeled EVs were recovered and used for the experiments.

To discard nonspecific fluorescent signal from antibodies, filtered PBS was incubated with each antibody separately and washed following the same protocol as the EVs incubated with the antibodies.

### EV lysis

As a control for specific EV detection, EVs were lysed by incubating with RIPA (1x; Merck Millipore) buffer for 30 min on ice, with periodic mixing every 5 min, as previously published (16).

### Flow cytometer specifications and setup

EV analysis, characterization, and sorting by flow cytometry were conducted using two distinct cell sorting platforms: the BD Influx™ cell sorter (Becton Dickinson, San José, CA) and the BD FACSDiscover™ S8 system equipped with CellView™ imaging technology (Becton Dickinson, San José, CA).

The BD Influx™ is a highly modular jet-in-air and customizable flow cytometer with low-pressure fluidics system allowing handling of fragile particles and supporting high-speed, multi-parameter sorting with user-defined optical configurations. This instrument is equipped with 5 lasers and 14 detectors for fluorescence with 2 scatter detectors. UV laser (355nm) with 2 channels for fluorescence; Deep blue laser (457nm) with 2 channels for fluorescence; Blue laser (488nm) with 4 channels for fluorescence and scatter detectors (FSC and SSC); Yellow-Green laser (561nm) with 3 channels for fluorescence; Red laser (640nm) with 3 channels for fluorescence. Data acquisition, instrument control, and cell sorting were performed using BD FACS™ Sortware.

The BD FACSDiscover™ S8 incorporates next-generation features, combining full-spectrum flow cytometry with real-time imaging through BD CellView™ technology. This integration facilitates the discrimination and sorting of small particles on both fluorescence emission and particle visualization, thereby expanding analytical capabilities for high-dimensional, image-assisted single-particle analysis. This instrument is equipped with 5 lasers and 78 detectors for fluorescence with 3 scatter detectors. Additionally, 6 detectors (3 for scatter and 3 for image fluorescence) related to the image module. UV laser (349nm) with 22 channels for fluorescence; Violet laser (405nm) with 20 channels for fluorescence and SSC detector; Blue laser (488nm) with 16 channels for fluorescence and scatter detectors (FSC and SSC); Yellow-Green laser (561nm) with 12 channels for fluorescence; Red laser (640nm) with 8 channels for fluorescence. Additionally, this technology incorporates de imaging module in the blue laser (488) 3 scatter detectors for imaging and 3 ones for image fluorescence. Data acquisition, instrument control, and cell sorting were performed using BD SpectralFX™ and BD CellView™ Image Technologies softwares.

Before each experimental session, both the conventional flow cytometer and the spectral flow cytometer underwent the quality control and calibration procedures recommended by the respective manufacturers to verify the correct performance of the optical system, lasers and detectors. Instrument performance was assessed using the corresponding commercial quality-control reference beads before sample acquisition, following current recommendations for standardized extracellular vesicle flow cytometry and reporting (13,17,18).

BD Influx™ cell sorter laser alignment and performance were carried out on a daily base using standard 8-peak rainbow calibration beads (Sphero™ Rainbow Calibration beads; BD Biosciences, Cat. No. 559123), following routine procedures for laser alignment. Drop delay calibration for sorting accuracy was carried out using recommended procedure with BD FACS™ Accudrop Beads (BD Biosciences, Cat. No 345249). All parameters were and analyzed using BD FACS Sortware™, version 1.0.0.0.650 (Becton Dickinson, San Jose, CA).

BD FACSDiscover™ S8 laser alignment and instrument performance were carried out on daily base using BD FACSDiscover™ Setup Beads (BD Biosciences, Cat. No. 665056). Drop delay calibration for sorting accuracy was carried out using recommended procedure with BD FACS™ Accudrop Beads (BD Biosciences, Cat. No 345249). Additionally, Image module was also calibrated using BD Cellview^TM^ Calibration beads (BD Biosciences, Cat. No. 665055). All parameters were acquired and analyzed using BD FACSChorus Software, version 6.2 (Becton Dickinson, San Jose, CA).

### NIST beads preparation, size scatter calibration

To characterize the sensitivity of both platforms for the detection of nanosized particles, an optical calibration based on light scatter was subsequently performed using National Institute of Standards and Technology (NIST)-traceable polystyrene beads (Thermo Fisher Scientific; catalogue numbers 3070A, 3080A, 3090A, 3100A, 3125A, 3150A, 3200A, 3240A, 3300A and 3400A), covering a certified diameter range between 70 and 400 nm. Serial dilutions of each bead population were prepared in phosphate-buffered saline (PBS) following the FCM_PASS_ protocol (19). Briefly, bead stocks were initially diluted 1:50, followed by working dilutions of 1:10, 1:100 and 1:1000. The optimal acquisition concentration corresponded to an approximate final dilution of 1:5000 for all bead populations except the 70-nm beads, which were acquired at approximately 1:500 to maximize the signal-to-background ratio.

Beads were acquired using the lowest available flow rate to minimize the event coincidence and swarming, during 60 s or until bead population was clearly visible from noise. During this calibration step, the acquisition threshold was established on the side scatter (SSC) parameter. In the conventional cytometer, SSC was generated by a single high-powered blue laser (Coherent 488 nm, 200 mW), whereas in the spectral flow cytometer SSC from the 405-nm violet laser was selected because of its improved sensitivity for the detection of small particles.

Light scatter calibration was performed using FCM_PASS_, which models light-scatter intensity according to Mie theory (19) while considering the refractive index of the calibration particles together with the optical configuration of each instrument. This approach generated an instrument-specific relationship between SSC intensity and equivalent scattering diameter, allowing the operational detection range of each cytometer to be standardized.

Following light scatter calibration, the acquisition threshold was shifted from SSC to the fluorescence channel while maintaining all remaining instrumental parameters unchanged, including detector gains and SSC settings. This strategy preserved the previously established SSC calibration while substantially improving the sensitivity for the detection of fluorescently labelled extracellular vesicles by minimizing electronic background noise.

Equivalent particle diameter was estimated from the calibration curve generated with FCM_PASS_ and used exclusively as a standardized optical parameter to characterize EV populations. Particle identification relied primarily on fluorescence rather than scatter intensity, thereby minimizing potential biases associated with differences in refractive index between biological vesicles and synthetic calibration beads.

### Fluorescence calibration and acquisition settings

To determine the optimal instrument configuration, a panel of yellow, fluorescent polystyrene nanobeads with defined diameters (1.34 μm, 0.88 μm, 0.45 μm, 0.22 μm, and 0.13 μm; Sphero™ Nano Polystyrene Yellow Beads, Spherotech refs NFPPS-0152-5, NFPPS-0252-5, NFPPS-0552-5, NFPPS-0852-5 and NFPPS-1552-5) and Rosetta Calibration Beads (Exometry B.V., Amsterdam, The Netherlands) were employed to systematically evaluate instrument performance for nanoparticles as well as sorting configuration (nozzle size, sheath pressure, and piezoelectric drive frequency). Based on these assessments, a 100 μm nozzle was selected, as it permits operation at relatively low sheath pressures (20 psi), thereby reducing particle velocity and increasing the transit time through the laser beam, an important parameter for enhancing time-of-flight and signal resolution.

### Extracellular vesicle acquisition and immunophenotyping on flow cytometers

Extracellular vesicle samples were acquired on both the conventional and spectral flow cytometers with fluorescence serving as the trigger parameter and SSC recorded as an additional characterization parameter. This acquisition strategy maximized the sensitivity for detecting labelled EVs while preserving the scatter information required for size standardization.

Immunophenotypic characterization of extracellular vesicles was performed using fluorochrome-conjugated antibodies directed against specific surface markers (see EV labelling section). Individual EV subpopulations were identified according to their fluorescence profiles and, when required, were subsequently isolated by fluorescence-activated sorting using the same acquisition strategy.

For the BD Influx^TM^ measurements acquisition was triggered on green fluorescence (FITC channel at 530/40 nm) and CFSE fluorescence was collected in the same FITC channel 530/40 nm. For BD FACSdiscover^TM^ S8 experiments, only CFSE-stained EVs and protein-enriched SEC fractions were assessed. Acquisition was also triggered on green fluorescence (B2 (515) channel at 517/16 nm), and CFSE fluorescence was detected in the same B2 (515) channel at 517/16 nm.

In both cytometers each sample was acquired for 60 s and event counts for each acquisition were recorded and are displayed within the corresponding figures in the results section.

Previous to running the experiments several control samples were analyzed: 0.22 µm-filtered Mili-Q water, 0.22 µm-filtered PBS, Unstained PPP-EV or iWJ-MSC-EV, CFSE-stained cell-free conditioned medium, CFSE-stained protein-enriched fraction from SEC, PBS with antibodies and CFSE-stained unlabeled PPP-EV or iWJ-MSC-EV.

### Fluorescence-activated EV sorting

EVs isolated from MSC-conditioned medium and PPP were first stained with CFSE and subsequently labelled with fluorophore-conjugated antibodies: anti-CD44-PE-Cy5 for MSC-EVs and anti-CD41-PE for PPP-EVs, as explained above. After antibody washing, equal volumes (1:1) of MSC-EVs and PPP-EVs were mixed together to obtain the final sorting mix. A comprehensive panel of controls was prepared in parallel: (i) unstained EV, (ii) CFSE-only stained EVs, (iii) CFSE-stained, single-antibody labelled EVs (CFSE^+^CD41-PE or CFSE^+^CD44-PE-Cy5), (iv) antibody-only controls prepared in filtered PBS (SEC elution buffer), (v) CFSE-stained EV lysed with RIPA, and (vi) CFSE-stained concentrated cell-free medium passed through a SEC column. These controls were used to establish fluorescence thresholds, validate specificity of staining and gating.

Flow cytometric analysis and sorting were performed on two platforms: the BD Influx^TM^ and the BD FACSdiscover™ S8. Instrument alignment and nanosized particle sensitivity were verified before each experiment as explained in the “Cell Sorter setup optimization” section. Sorting gates were defined as P1 (CFSE⁺ EVs), P2 (CD41-PE⁺ PPP-EVs), and P3 (CD44-PE-Cy5⁺ MSC-EVs). Neither color compensation on the BD Influx^TM^ nor spectral unmixing on the BD FACSdiscover™ S8 was required to manage fluorescence spillover.

Post-sort purity was determined by analysing each sorted fraction using identical gating strategies and was expressed as the proportion of CFSE^+^CD41^+^ or CFSE^+^CD44^+^ events relative to total events.

Sorting performance varied across instruments. On the BD-Influx™, sorting runs lasted 3-5 h without achieving complete recovery. By contrast, the BD FACSdiscover™ S8 completed equivalent sorts within 1-2 h to reach equivalent yield, showing better sorting efficiency (Fig. 1).

**Figure 1.**
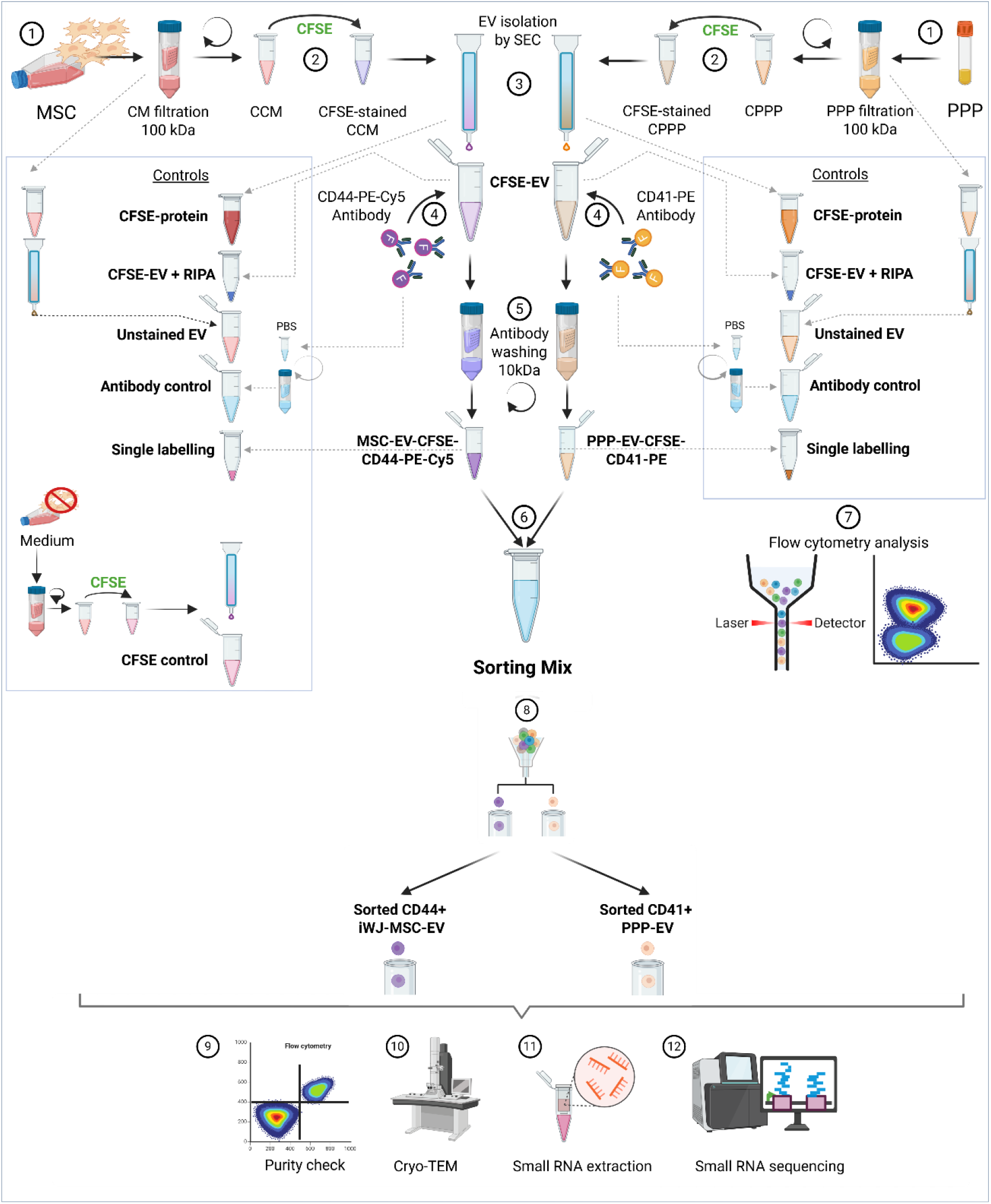
Nanoparticle Fluorescence-activated vesicle sorting (NanoFACS) workflow. Overview of the experimental pipeline from EV preparation to downstream analysis. (1) Collection and concentration of mesenchymal stromal cell (MSC)–conditioned medium (CM) and platelet-poor plasma (PPP). (2) CFSE staining of concentrated CM (CCM) and PPP. (3) EV isolation from stained samples by size exclusion chromatography (SEC). (4) Antibody labelling of EVs with CD44-PE-Cy5 (MSC-EVs) or CD41-PE (PPP-EVs). (5) Removal of unbound antibodies using 10 kDa filters. (6) Preparation of the sorting mix by combining both CFSE- and antibody-labelled EV populations at a 1:1 ratio. (7) Flow cytometry analysis of the sorting mix and all associated controls. (8) Sorting of CFSE⁺CD44⁺ MSC-EVs and CFSE⁺CD41⁺ PPP-EVs. (9) Purity check. (10) Cryo-transmission electron microscopy (Cryo-TEM) analysis of pre- and post-sort fractions. (11) RNA extraction from pre- and post-sort fractions. (12) Small RNA sequencing of pre- and post-sort samples.

### Cryogenic transmission electron microscopy (Cryo-TEM)

To confirm EV presence and their integrity, post-sort PPP-EVs and iWJ-MSC-EVs were analyzed by cryogenic-electron microscopy (cryo-TEM) using a transmission electron microscope (Joel JEM 2011, Tokyo, Japan), as previously described (20). Before the analysis, samples were concentrated with an Amicon Ultra filter of 10 KDa (Millipore, Billerica, MA, USA) at 3000 ×g for 10 min. For each sample, minimum 10 pictures from representative fields were obtained at different magnification scales (1000 nm, 500 nm, and 200 nm).

### RNA extraction and quantification

The RNA from the pre-sort and post-sort samples was obtained using the miRNeasy Micro Kit (Qiagen). Briefly, samples were concentrated using a 2 mL Amicon Ultra filter of 10 KDa (Millipore, Billerica, MA, USA) at 3000 ×g for 10 min. Then, the concentrate was mixed 1:1 with Qiazol and the remaining protocol was performed following manufacturer’s instructions.

### RNA sequencing

Before starting with the smallRNAseq experiments all samples were quantified and measured using the Bioanalyzer RNA 6000 pico assay at the 2100 Bioanalyzer System (Agilent) that provides reliable and reproducible characterization of RNA from multiple sample types, from as little as 50 pg. To prepare the small RNA libraries, the NEXTFLEX Small RNA-Seq Kit v4 for Illumina Platforms (Revvity) was performed. The protocol comprises ligation of both 3’ and 5’ adapters to the RNA molecules, reverse transcription to generate cDNA, and amplification and indexing by PCR of cDNA molecules that have adapters on both ends. PCR is performed using the UDI barcoded primer mix included in the kit, so final libraries have unique dual indexes. Final libraries were analyzed using Agilent Bioanalyzer HS DNA Assay (Agilent) and quantified by qPCR using the KAPA Library Quantification Kit (Roche) to check the measurement of the library insert and its concentration. Adapters with different indexes were used to be able to sequence several samples in a single lane. All samples were sequenced using the Nextseq2000 Illumina sequencer at a sequencing depth of 20 M SE 1x50bp reads per sample. Following Illumina’s quality specifications, more than 80% of the bases that passed the quality filters had a quality value greater than Q30 (Phred score) and 90% >Q20.

### RNAseq data analysis

Raw sequencing reads were processed to remove adapter sequences and low-quality bases using Skewer version 0.2.2 (21). Reads shorter than 19 nucleotides after trimming were discarded. The remaining reads were aligned to the human genome (hg38) using ShortStack version 3.8.5 (22). Annotation of aligned reads was performed against genecode version 37. A count matrix was subsequently generated, and genes with a total read count of fewer than 10 across all samples were excluded from further analysis.

Differential expression analysis between sorted fractions was carried out using the DESeq2 package (23) (version 1.34.0) in R, applying a false discovery rate (FDR) correction for multiple testing (adjusted p-value >0.05).

For data visualization, principal component analysis (PCA) was performed to assess clustering between pre- and post-sort PPP-EV and MSC-EV fractions. A volcano plot was generated to display differentially expressed small RNAs between the post-sort PPP-EV and MSC-EV populations. Both PCA and volcano plot representations were produced using GraphPad Prism v.10 (GraphPad Software, USA).

## Results

### Instrument calibration reveals enhanced scatter sensitivity of spectral flow cytometry

To establish the analytical sensitivity of both flow cytometry platforms for nanosized particle detection, conventional and spectral flow cytometers were initially calibrated using NIST-traceable polystyrene beads ranging from 70 to 400 nm in diameter. Calibration was performed using a side scatter (SSC)-based acquisition threshold, allowing the optical response of each instrument to be characterized under standardized conditions.

The selection of an appropriate trigger parameter is critical for the detection of nanoparticles, and has been widely debated in the literature, particularly regarding the suitability of scatter parameters such as side scatter (SSC), or fluorescence-based triggering strategies (10,24). In the conventional platform, SSC was detected with the blue laser together with a PMT (Photomultiplier Tube) detector, while we applied the use of violet-laser SSC detection together with APD detectors (Avalanche Photodiodes) in the spectral cytometer, which improves light scatter resolution for nanosized particles.

Under these acquisition settings, the conventional flow cytometer consistently failed to resolve the smallest bead populations above buffer background noise with particles below approximately 200 nm (Fig. 2A and B) becoming progressively indistinguishable from background noise (data not shown). The spectral flow cytometer successfully resolved all bead populations across the complete calibration range, including the 70-nm reference particles (Fig. 2C and D), consistent with the increased scattering efficiency achieved using shorter excitation wavelengths (25).

**Figure 2.**
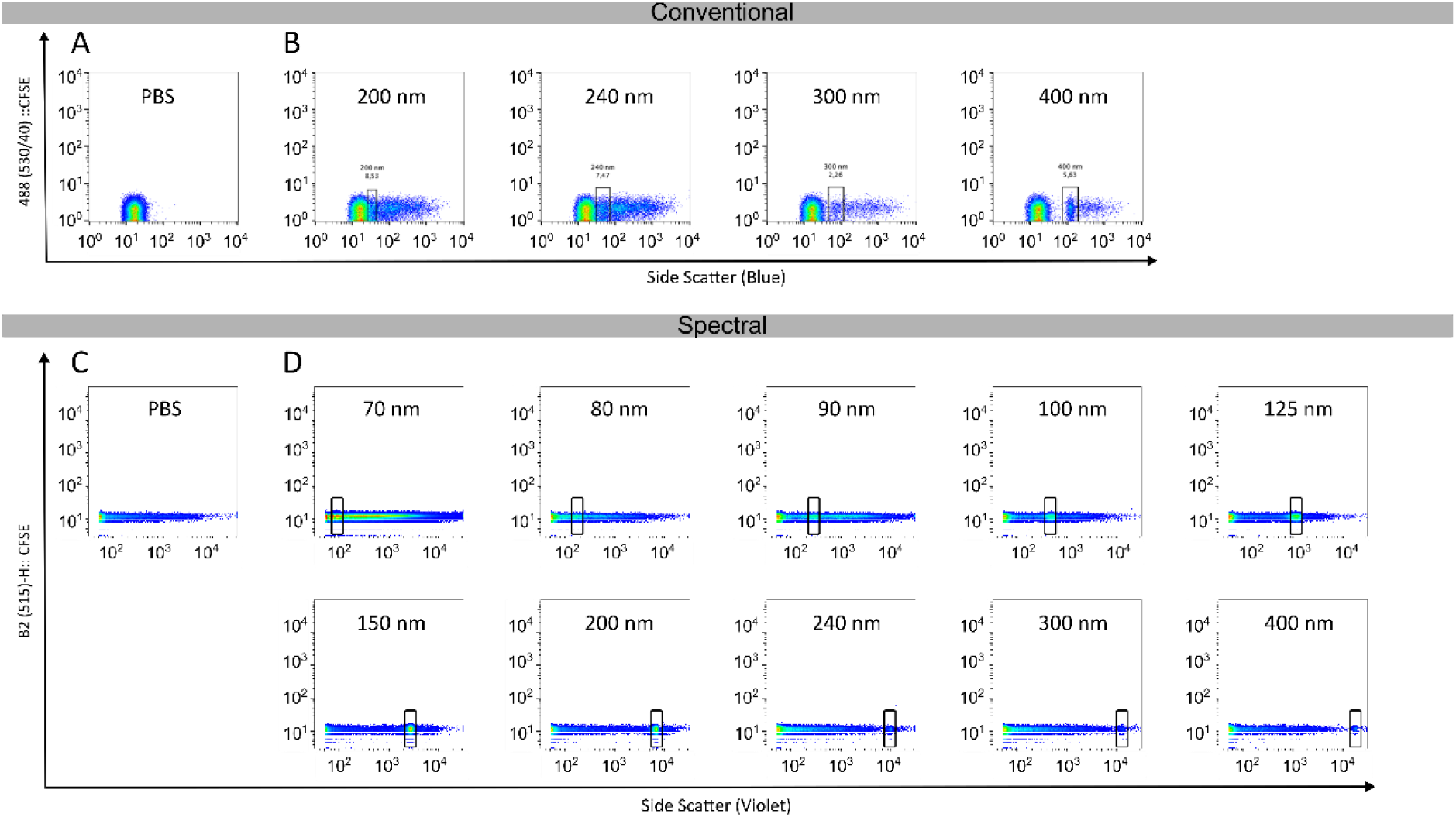
Spectral flow cytometry better resolves NIST-traceable polystyrene beads using scatter-based calibration than conventional flow cytometry platforms. Representative SSC plots obtained using the BD Influx conventional flow cytometer (A-B) and the BD S8 spectral flow cytometer (C-D). (A) PBS background acquired on the conventional cytometer. (B) NIST-traceable polystyrene bead mixture containing particles ranging from 200 to 400 nm acquired on the conventional cytometer. Smaller-sized beads were not discriminated from background noise and are not shown. (C) PBS background acquired on the spectral cytometer. (D) NIST-traceable polystyrene bead mixture containing particles ranging from 70 to 400 nm acquired on the spectral cytometer. The spectral cytometer resolved bead populations across the complete size range, including the 70-nm population, whereas the conventional showed progressively reduced discrimination of the smallest bead populations from background noise.

These results demonstrate that spectral flow cytometry provides improved optical sensitivity during scatter-based calibration, particularly for particles approaching the lower detection limit of conventional flow cytometers.

### Fluorescence-triggered acquisition preserves scatter calibration while improving EV detection

Following scatter calibration, acquisition conditions were modified to increase sensitivity. Specifically, the acquisition threshold was transferred from SSC to the fluorescence channel, whereas all remaining instrumental settings, including SSC detector gain and optical configuration, were maintained unchanged.

Therefore, we implemented a fluorescence-based triggering strategy. The fluorescence signal was directed through a 505 nm long-pass (LP) filter and collected using a 530/40 nm band-pass filter on the BD Influx™ and through B2 (515) channel at 517/16 nm on the BD FACSdiscover^TM^ S8.

Under fluorescence-triggered acquisition, both cytometers detected all fluorescent reference particles included within the calibration range (Fig. 3). We successfully detected the fluorescent Spherotech yellow nanobeads (Fig. 3A and B) and the Rosetta calibration beads, which contain a mixture of seven types of polystyrene beads with diameters between 100 nm and 1000 nm, and one-sized fluorescent beads of 100 nm (Fig. 3C and D). Because these particles incorporate intrinsic fluorescence, they more closely reproduce the optical conditions encountered during EV acquisition than non-fluorescent polystyrene beads. Their detection therefore provided an additional indication that the selected acquisition strategy would be appropriate for the expected experimental conditions.

**Figure 3.**
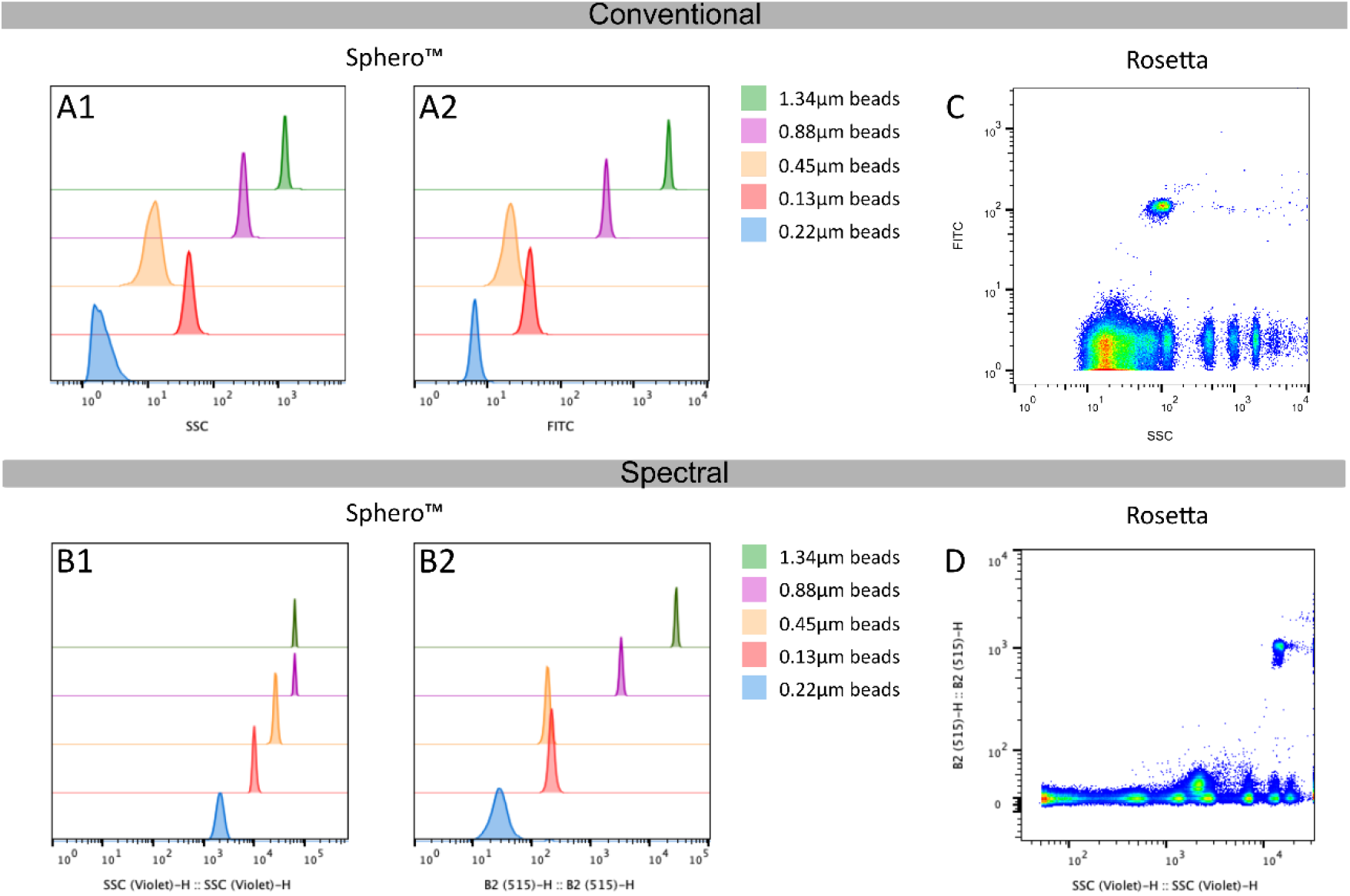
Fluorescence-triggered detection of fluorescent reference particles preserves side scatter sensitivity. Fluorescent reference beads were acquired using the same fluorescence-triggered settings subsequently applied for extracellular vesicle analysis. (A) Intensity histograms for SSC (A1) and the FITC channel (A2) of Spherotech yellow fluorescent nanobeads (0.13, 0.22, 0.45, 0.88, and 1.34 µm) acquired on the conventional BD Influx flow cytometer. (B) Intensity histograms for SSC (B1) and FITC in the B2 channel (B2) of the same Spherotech bead populations acquired on the BD S8 spectral flow cytometer. (C) Rosetta calibration beads acquired on the BD Influx under fluorescence-triggered acquisition. (D) Rosetta calibration beads acquired on the BD S8 under fluorescence-triggered acquisition. Detection of the fluorescent bead populations on both platforms confirmed that fluorescence-triggered acquisition was compatible with the scatter calibration established during instrument standardization and with the acquisition conditions used for EV analysis

Collectively, these observations demonstrate that fluorescence-triggered acquisition substantially improves the sensitivity for fluorescent nanoparticle detection while preserving the scatter resolution established during instrument calibration.

### Fluorescence-triggered acquisition enables robust identification of small EVs

Fluorescently labelled EVs derived from immortalized Wharton’s Jelly-derived mesenchymal stromal cells (iWJ-MSCs) isolated by size exclusion chromatography (SEC) were subsequently acquired using the fluorescence-triggered strategy established during calibration. Under these conditions, both the conventional and spectral flow cytometers clearly identified the EV population above the background events generated by electronic noise or non-specific scatter (Suppl. Fig. 1).

Conversion of SSC intensity into equivalent scattering diameter using the FCM_PASS_ calibration model indicated that the majority of detected vesicles were distributed between an equivalent diameter of approximately 90 to 200 nm (EV population being between the 90 nm NIST beads in turquoise and the 200 nm-NIST beads in light green in Fig. 4A and B). The model was consistently replicated using the Rosetta Beads instead of the NIST for calibration (Fig. 4C and D). It should be emphasized that these values represent equivalent optical diameters derived from light-scatter modelling and should therefore not be interpreted as direct measurements of vesicle size, but rather as standardized optical diameters derived from Mie modelling (19).

**Figure 4.**
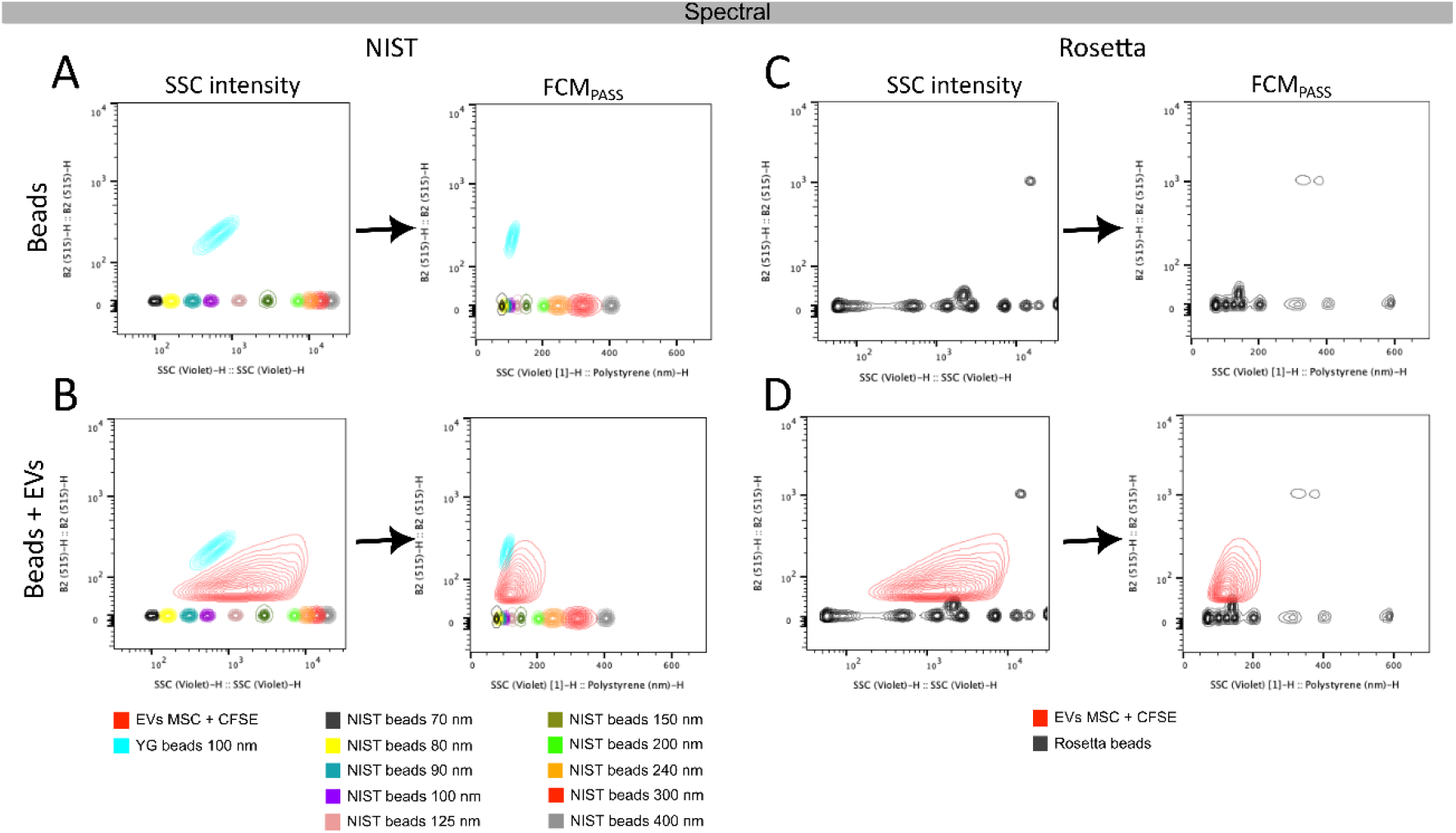
Fluorescence-triggered acquisition enables robust identification of small EVs. Scatter signals from CFSE-fluorescently labelled extracellular vesicles (EVs) were transformed to scattering diameter using two bead-based calibration approaches on the S8 spectral flow cytometer. (A) SSC and B2 channel intensity distributions of NIST-traceable polystyrene beads together with 100-nm YG beads (left) and the corresponding FCM_PASS_ transformation to diameter (nm; right). (B) SSC and B2 channel distribution of CFSE-stained EVs and the corresponding FCM_PASS_-derived equivalent scattering diameter using the NIST/YG calibration (right). (C) SSC and B2 channel intensity distributions of Rosetta calibration beads together with 100-nm YG beads (left) and the corresponding FCM_PASS_ transformation (right). (D) SSC and B2 channel distribution of CFSE-stained EVs (left) and its equivalent scattering diameter using the Rosetta/YG calibration (right).

Because the optical configuration of the BD Influx^TM^ conventional cytometer is not supported by the FCM_PASS_ calibration framework, conversion of scatter intensity into equivalent scattering diameter could only be performed for the BD FACSdiscover^TM^ S8 cytometer. However, the raw data acquired in the BD Influx^TM^ shows a similar pattern as the one observed in the spectral cytometer confirming the size of the EV between 100 nm and 200 nm either in the comparison with the NIST beads (Suppl. Fig. 1A and B) or the Rosetta Beads (Suppl. Fig. 1C and D).

Overall, fluorescence-triggered acquisition enabled robust identification of small EVs in both cytometry platforms while preserving standardized scatter characterization. However, the spectral cytometer generated a broader dynamic range of detectable scatter signals, enabling a more robust calibration of equivalent particle diameter using FCM_PASS_.

### Sample dilution and flow rate modulate swarming and differentially affect EV and protein fraction detection

To evaluate potential coincident particle detection (swarming) and determine the dilution range compatible with single-EV detection, ten serial two-fold dilutions of CFSE-labelled EV derived from iWJ-MSC (MSC-EV) or Platelet-Poor Plasma (PPP-EV) and their corresponding SEC protein fractions (MSC-Prot and PPP-Prot) were prepared in PBS and analyzed on both flow cytometry platforms. **Fig. 5** shows dilutions from 1:32 onwards, corresponding to the range in which EV fractions started to display an approximately linear decrease in the detected event number.

**Figure 5.**
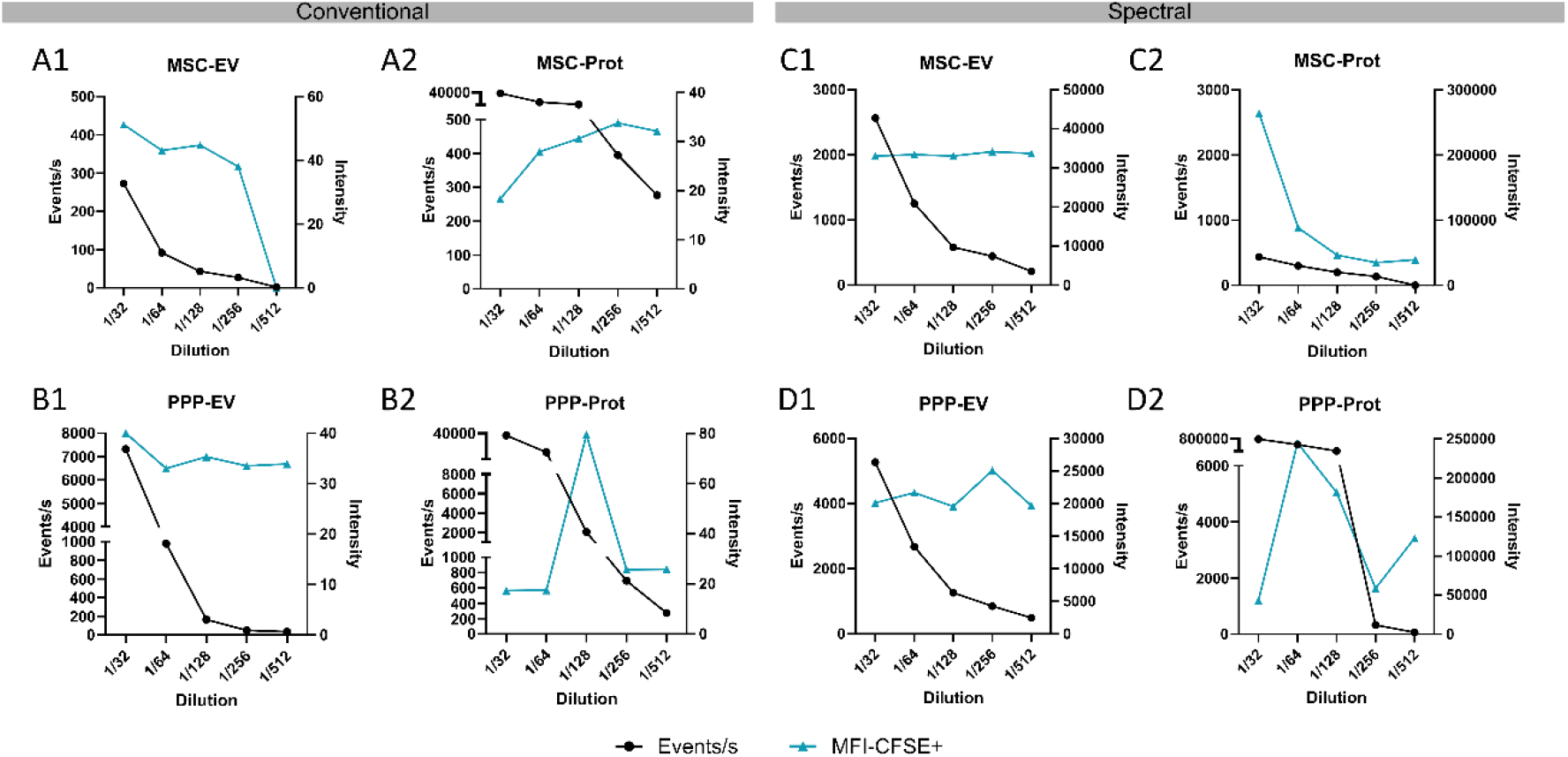
Effect of sample dilution on swarming and EV or protein fraction detection. Two-fold serial dilutions of CFSE-labelled EV and protein SEC fractions from MSC or PPP MSC were analysed from the 1:32 to 1:512 dilution, evaluating the number of CFSE^+^ events (60 sec) and the CFSE median fluorescence intensity (MFI) for each sample. (A) MSC-derived and CFSE-labelled EV (A1) and Protein fraction (A2) acquired on the conventional BD Influx flow cytometer. (B) PPP-derived and CFSE-labelled EV (B1) and Protein (B2) acquired on the BD Influx. (C) MSC-derived and CFSE-labelled EV (C1) and Protein (C2) acquired on the spectral BD S8 flow cytometer. (D) PPP-derived and CFSE-labelled EV (D1) and Protein (D2) acquired on the BD S8. EV fractions showed an approximately linear decrease in event number with relatively stable MFI, whereas protein fractions showed high event counts at lower dilutions and greater variation in MFI.

For both MSC-derived (Fig. 5 A1 and C1) and PPP-derived EVs (Fig. 5 B1 and D1), increasing dilution resulted in a progressive and approximately linear reduction in event counts, while median fluorescence intensity (MFI) remained relatively stable across the analyzed dilution range. Of note, the linearity of the event counts within the working range of the spectral flow cytometer was better correlating to the experimental dilution than the conventional flow cytometer.

In contrast, protein fractions (Fig 5. A2, B2, C2, and D2) showed substantially higher event numbers at lower dilutions and rapidly reached saturation. When a linear decrease in event number was observed in protein fractions, this was accompanied by marked changes in MFI. These results indicate a clear difference in dilution behavior between EV and protein fractions. The high number of protein-associated events observed at lower dilutions would hinder discrimination of EV-derived events in non-separated samples, supporting the need for prior EV isolation before flow cytometric detection and sorting.

Similar to the dilution factor, another key variable influencing the potential “swarming effect”, the accuracy and reproducibility of EV detection by flow cytometry is the flow rate; the speed at which the sample passes through the interrogation point. To assess its own background signal and EV detectability, we analyzed CFSE-labeled iWJ-MSC-EVs alongside several control samples, including sterile-filtered water, CFSE-labeled protein-enriched SEC fractions, and CFSE-labeled medium processed in the absence of cells (Suppl. Fig. 2). For the conventional cytometer samples were acquired at increasing flow rates corresponding to sheath pressures of 21.4, 21.5, 21.8, and 25 psi. Across all control samples, a progressive increase in flow rate led to a corresponding rise in CFSE⁺ events (Suppl. Fig. 2A–C), consistent with a flow rate-dependent background artifact. This effect was most pronounced in the protein-enriched fraction, followed by the medium-only control, and was least evident in the water control, reflecting differences in free dye or protein aggregate content.

In contrast, the CFSE-labeled iWJ-MSC-EV sample exhibited an opposing trend: increasing flow rates were associated with a gradual decrease in the CFSE⁺ percentage (Suppl. Fig. 2D). This suggests that higher flow rates reduce the cytometer’s ability to resolve small, low-intensity particles such as EVs, likely due to shorter dwell times and diminished signal integration.

To determine whether flow rate similarly affected EV detection under spectral flow cytometry conditions, we next evaluated increasing flow rates of 1, 20 and 50 relative flow rates, corresponding to 0.16, 0.31 and 0.75 μL/s respectively, using CFSE-labeled MSC-EVs and the corresponding protein-enriched fraction (Suppl. Fig. 2E and F). For CFSE-labeled EVs, the signal remained relatively stable at low and moderate flow rates but became indistinguishable from background at the highest flow rate, where the population appeared broadly CFSE⁺, indicating loss of resolution (Suppl. Fig. 2E). The protein fraction also showed a marked flow rate-dependent response, with artefactual CFSE⁺ detection at the highest flow rate, similarly resulting in loss of discrimination (Suppl. Fig. 2F). Together, these data underscore the importance of optimizing flow rate parameters to minimize false-positive readouts and preserve accurate detection of true EV events and show how the analysis of the right control samples can help set it up.

### Detection of nanosized Extracellular Vesicles by Flow cytometry

To assess the capability of the conventional flow cytometer to detect nanoscale particles, we first determined the instrument’s sensitivity by assessing the background signal of electronic noise and the one generated by 0.22 µm-filtered water (Fig. 6A and B). Next, we proceeded to analyze MSC-EVs isolated by SEC. Initial analysis of unstained EV samples revealed no discernible signal above the electronic noise threshold, indicating the need for fluorescent staining to detect individual vesicles (Fig. 6C). For this, iWJ-MSC-derived EVs were fluorescently labeled with CFSE and purified via SEC to remove unbound dye at the same time of EV isolation (Fig. 6D), as described in the Methods section. Bead-assisted flow cytometry corroborated CFSE staining of EV, peaking at Fractions 4-6, coinciding with CD63^+^CD90^+^, but depleted of free protein present in the preparation, which eluted later in F7-onwards and was also stained with CFSE. Nevertheless, only CFSE-labeled EV were detectable above noise by BD Influx (Fig. 6E), with CFSE^+^ events dramatically decreasing in fractions F7-onwards, consistent with the decline of EV presence in those fractions, and corroborating an EV-associated signal (Fig. 6E and F). Of note, the CFSE-EV cloud was overlapping with the background, indicating that EV signal is within the detection limit of the machine (Fig. 6E). To ensure the true gating of EV as CFSE^+^ events, the CFSE-labeled protein-enriched fractions from the same preparation were analyzed as a control (Fig. 6G). The CFSE-labeled protein pool did not generate a comparable positive signal as CFSE^+^EV and showed instead the same CFSE threshold as unstained EVs (Fig. 6G). To control for potential non-specific CFSE staining or detection, culture medium alone was subjected to the same concentration, staining, and SEC purification procedures. The corresponding EV-like elution fractions from this cell-free control displayed only background fluorescence signal, indicating dismissible unspecific detection of medium-derived events (Fig. 6H, right).

**Figure 6:**
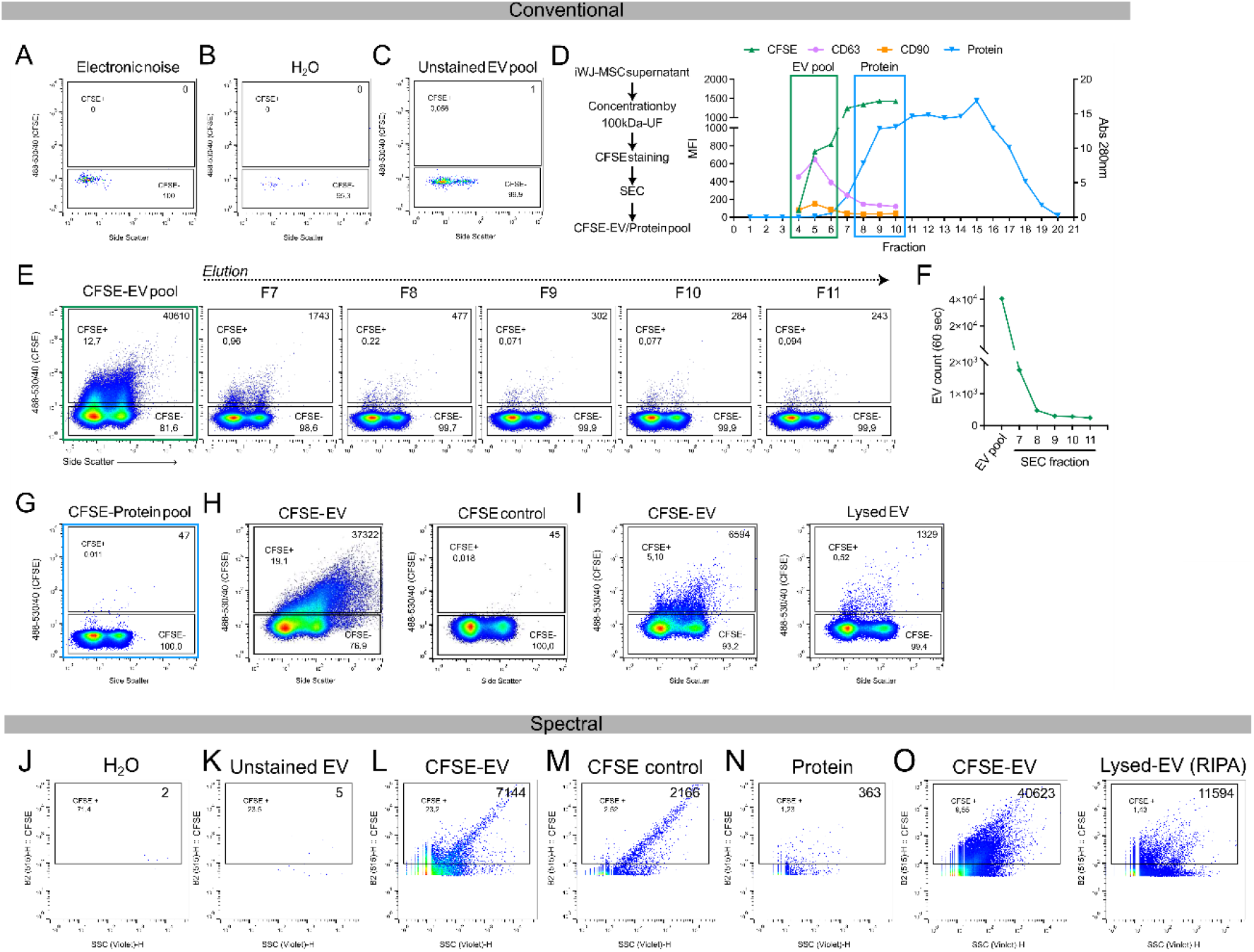
Evaluation of EV detection by conventional (A-I) and spectral (J-O) flow cytometry. (A, B, J, K) Background signal for (A) electronic noise, (B, J) 0.22 µm-filtered water and (C, K) unstained iWJ-MSC-EVs. (D) Schematic of iWJ-MSC supernatant processing for CFSE-labeled EV obtention (left) and representative size-exclusion chromatography (SEC) profile of CFSE-labeled concentrated supernatant from iWJ-MSC. CFSE⁺ iWJ-MSC-EV-containing fractions are identified as positive for CD63 and CD90 markers (right). The EV- and protein-pooled fractions are framed in green and blue, respectively. (E) Conventional flow cytometric analyses of CFSE-labeled iWJ-MSC-EVs and its following SEC fractions. (F) Events count of the CFSE⁺ gate for the EV-pool and the following SEC fractions, corresponding to (E), for 60 seconds at a constant flow rate. (G-I) Representative plots of the CFSE-labeled protein pool (G), CFSE-labeled EVs and CFSE-labeled cell-free basal medium control (H), CFSE-labeled EVs before and after RIPA buffer lysis (I), to evaluate signal specificity for intact vesicular structures. (L-N) Spectral flow cytometric analysis of CFSE-labeled iWJ-MSC-EVs (L), CFSE-labelled cell-free conditioned medium processed by SEC, pooled for EV-eluting fractions (M), iWJ-MSC-derived protein-enriched SEC fraction (N). (O) Assessment of signal specificity through lysis of CFSE-labeled EVs. The CFSE⁺ count (60 sec) is indicated in the top right corner of each plot. Data representative of 2-3 independent experiments.

Finally, to further confirm the vesicular nature of the CFSE-labeled detected events, we lysed the labeled EV sample with RIPA buffer. This treatment, known to destroy intact EV (16), substantially reduced the positive fluorescent signal 5-fold (Fig. 6I), supporting the conclusion that the CFSE-positive events detected by flow cytometry corresponded to intact membrane-bound vesicles rather than dye aggregates or protein complexes.

To determine whether the same fluorescence-based detection strategy could be applied to the spectral flow cytometer, we evaluated the equivalent set of controls used for the conventional platform. Background signal was first assessed by analyzing sterile-filtered water and unstained (CFSE⁻) iWJ-MSC-derived EVs (Fig. 6J and K). Due to the adjustable lower limit of the fluorescence threshold in the BD FACSdiscover^TM^ S8, virtually no background signal was detected in any of the control samples. We then analyzed CFSE-labeled MSC-EVs, which produced a clear CFSE⁺ signal above the background (CFSE⁻) threshold (Fig. 6L). This signal was markedly reduced when analyzing CFSE-labeled cell-free medium (Fig. 6M) or the protein-enriched SEC fraction (Fig. 6N), yielding only 2.52% (2,165 events) and 1.23% (363 events) CFSE⁺ events, respectively, compared to 23.2% (7,144 events) in the labeled EV sample. Similarly, lysis of CFSE-labeled EVs led to a reduction in CFSE⁺ events from the 9.55% (40,623 events) to 1.43% (11,594 events), confirming that the CFSE signal was associated with intact vesicular structures (Fig. 6O).

### EVs can be sorted by flow cytometry

We next assessed the capability of conventional flow cytometry to sort EVs based on fluorescent triggering. To establish sorting gates, we first analyzed the instrument’s electronic noise profile to determine CFSE-negative and CFSE-positive regions (Fig. 7A). Using this gating strategy, we then analyzed a CFSE-labeled iWJ-MSC-EV preparation, confirming the presence of a distinct CFSE⁺ vesicle population suitable for sorting (Fig. 7B).

**Figure 7.**
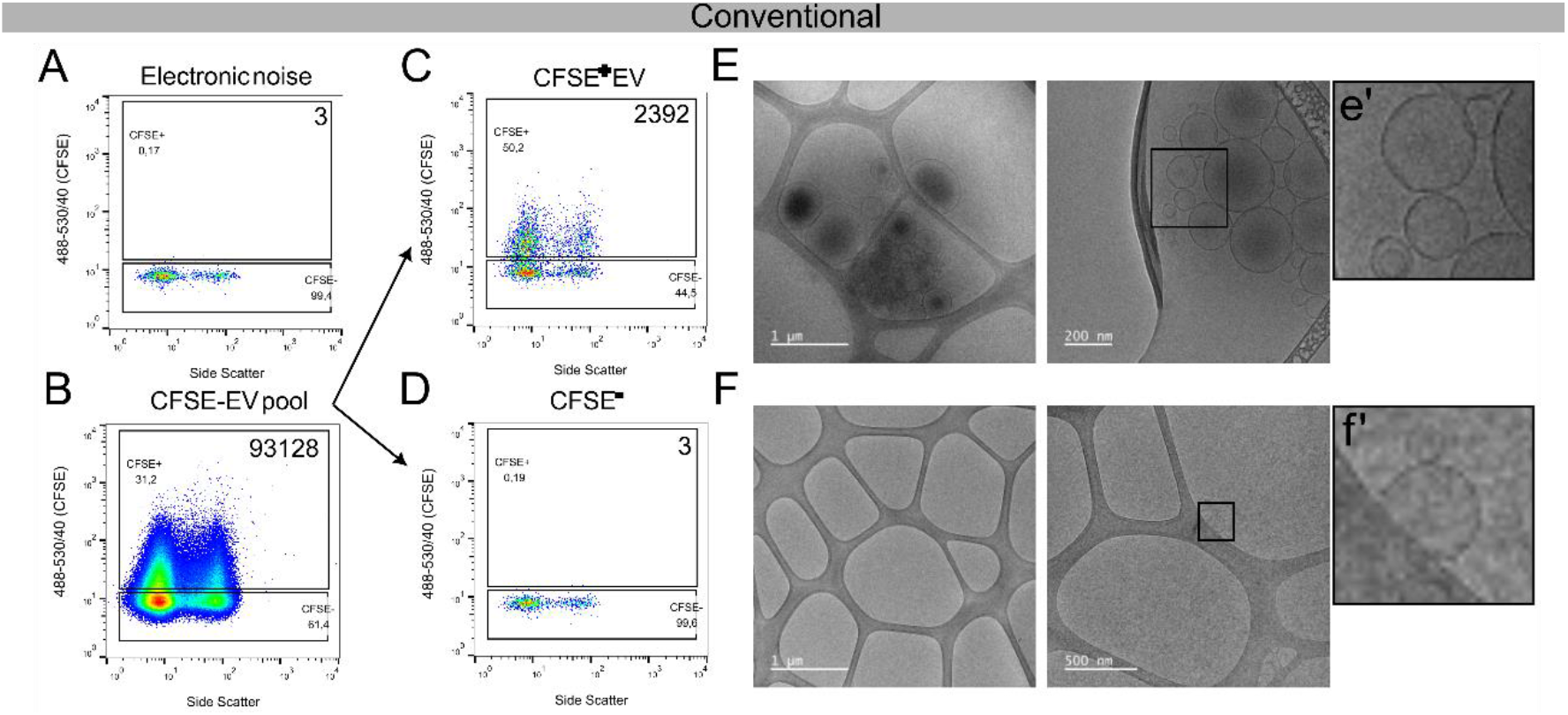
Sorting and characterization of CFSE-labeled EVs by conventional sorter. Representative plots of (A) the electronic noise background used to define the detection threshold, (B) CFSE-labeled iWJ-MSC-EV sample used for sorting, (C-D) post-sorting purity analysis of the CFSE⁺ (C) and CFSE^-^ (D) populations. (E-F) Cryo-electronic transmission microscopy (Cryo-TEM) pictures of the post-sort CFSE⁺ (E) and CFSE^-^ (F) sorted populations, showing nanosized, round-shaped EVs in the sorted CFSE^+^ events. A single vesicle was detected in the CFSE^-^ sorted fraction. Data representative of 21 cryo-TEM images analyzed. The CFSE⁺ events count during 60 s of acquisition are shown in the top right corner of each plot.

Fluorescence-based vesicle sorting was next performed to separate CFSE⁺ and CFSE⁻ particles. This procedure yielded two distinct fractions: a CFSE⁺ population with high purity, comprising 50.2% CFSE⁺ events and clearly distinguishable from background, indicating that the 44.5% in the negative fraction corresponds to noise (Fig. 7C), and a CFSE⁻ population with minimal CFSE⁺ contamination (0.19% CFSE⁺ events; Fig. 7D). To validate the vesicular nature of the sorted populations, both fractions were analyzed by cryogenic transmission electron microscopy (cryo-TEM). Vesicular structures consistent with the morphology and size of EVs were exclusively observed in the CFSE⁺ fraction (Fig. 7E), while the CFSE⁻ fraction showed an absence of detectable vesicles, except for a single event (Fig. 7F). The strong enrichment of vesicular structures in the CFSE⁺ fraction, as confirmed by cryo-TEM, and the absence of EV in the negative fraction demonstrated the feasibility of EV sorting by flow cytometry based on fluorescent triggering on CFSE staining.

### Antibody-labeled EV populations can be detected

To determine whether we could sort specific EV subtypes based on surface marker expression, we developed a new sorting strategy using antibody-labeled samples and appropriate controls. For this set up, a mixed EV sample consisting of CFSE-labeled PPP-EVs and iWJ-MSC-EVs was prepared. These EVs were labeled with lineage-specific anti-CD41-PE and anti-CD44-PE-Cy5 antibodies, respectively. A dedicated gating strategy was established based on the labeling and control conditions (Fig. 8A). First, CFSE⁺ events were selected (P1 gate), followed by the identification of CD41⁺ (P2 gate) and CD44⁺ populations (P3 gate) used for sorting. Unlabeled controls (no antibody labeling) revealed some non-specific PE signal in both PPP-EV and iWJ-MSC-EV samples, but more evident for PPP-EVs (Fig. 8B). Of note, fluorescence overlap compensation was discarded to avoid loss of signal. Instead, an additional gate (P4) was included to monitor PE-specific fluorescence, thereby helping to distinguish true CD41⁺ events from background.

**Figure 8.**
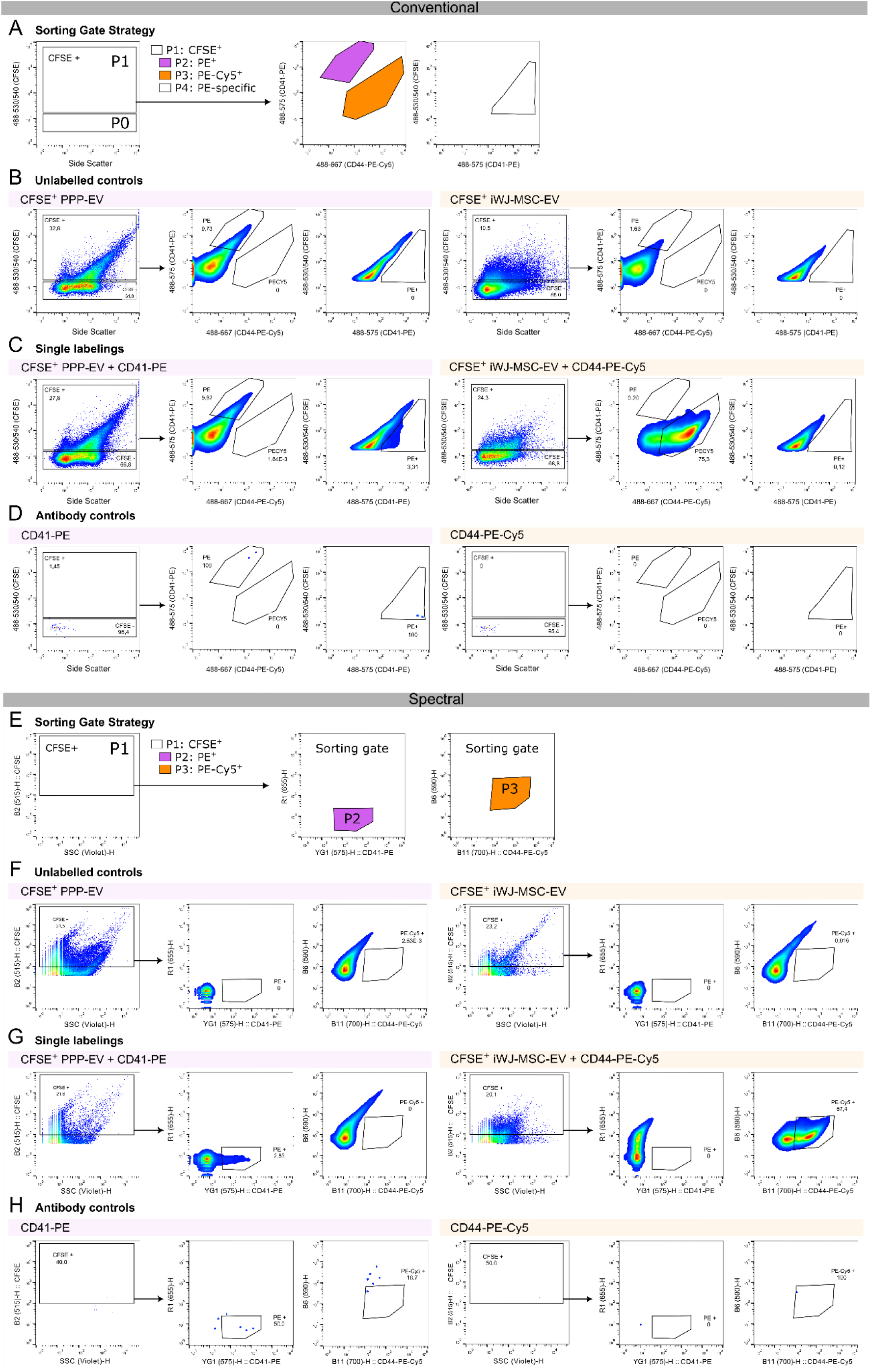
Evaluation of the conventional and spectral cell sorters for nanosized particle detection: gating strategy and controls for sorting antibody-labeled EVs. (A) Conventional flow cytometry gating strategy for EV sorting. P0: CFSE⁻ gate; P1: CFSE⁺ gate; P2: CD41-PE⁺ sorting gate; P3: CD44-PE-Cy5⁺ sorting gate; P4: CFSE⁺CD41-PE⁺ gate (PE specificity control). (B) Unlabeled controls: CFSE⁺ platelet-poor plasma-derived EVs (PPP-EVs) and CFSE⁺ iWJ-MSC-derived EVs. (C) Single-antibody labeling controls: CFSE⁺ PPP-EVs labeled with CD41-PE and CFSE⁺ MSC-EVs labeled with CD44-PE-Cy5. (D) Antibody-only controls: CD41-PE and CD44-PE-Cy5 incubated in EV elution buffer without sample, to assess background signal. (E) Spectral flow cytometry gating strategy for EV sorting. P1: CFSE⁺ gate; P2: CD41-PE⁺ sorting gate; P3: CD44-PE-Cy5⁺ sorting gate. (F) Unlabeled controls: CFSE⁺ platelet-poor plasma-derived EVs (PPP-EVs) and CFSE⁺ iWJ-MSC-derived EVs. (G) Single-antibody labeling controls: CFSE⁺ PPP-EVs labeled with CD41-PE and CFSE⁺ MSC-EVs labeled with CD44-PE-Cy5. (H) Antibody-only controls: CD41-PE and CD44-PE-Cy5 incubated in EV elution buffer without sample, to assess background signal.

Moreover, when each EV population was labeled with its corresponding antibody, this non-specific PE signal was no longer visible in the iWJ-MSC-EV sample (Fig. 8C) and the P4 gate proved sufficient to identify true CD41⁺ PPP-EVs. Clear populations of CD44⁺ events were also identified in the iWJ-MSC-EV sample, confirming the effectiveness of the antibody labeling. Moreover, antibody-only controls prepared in filtered PBS (the EV elution buffer) showed no detectable background signal (Fig. 8D), supporting the specificity of the labeling protocol.

While manual conventional cytometers are highly adaptable and versatile for tailored analysis like EV sorting, they are progressively being replaced by advanced, more sensitive spectral flow cytometers. Thus, we sought to replicate the setup and antibody-labeled EV sorting experiments using a spectral flow cytometer system.

To determine whether specific EV subtypes could also be sorted based on antibody labeling using the spectral cytometer, we replicated the conventional sorting strategy and established a corresponding gating approach tailored to the spectral Flow cytometer (Fig. 8E). Sorting gates were defined based on antibody labeling and control conditions: CFSE⁺ events were first selected (P1 gate), followed by identification of CD41⁺ (PE⁺, P2 gate) and CD44⁺ (PE-Cy5⁺, P3 gate) populations. Due to the improved resolution of the PE signal in the spectral cytometer, the yellow-green laser channel was used for PE detection. This adjustment eliminated non-specific PE signal in control samples, and thus no additional PE-specificity gate (as used in the BD Influx™) was required (Fig. 8F and 8G).

Sorting was performed using the P2 and P3 gates. Unlabeled controls (no antibody labeling) exhibited no background signal in either the PE (P2) or PE-Cy5 (P3) gates (Fig. 8F). In contrast, PPP-EVs and MSC-EVs labeled with the corresponding antibodies displayed distinct and specific signals within the expected gates (Fig. 8G). Furthermore, antibody-only controls prepared in filtered PBS (EV elution buffer) showed no detectable background signal (Fig. 8H), confirming the specificity and robustness of the antibody labeling protocol tested in the spectral cytometer.

### Antibody-labeled EV populations can be sorted

Next, the mixed sample of CFSE-labeled CD41-PE⁺ PPP-EVs and CD44-PE-Cy5⁺ iWJ-MSC-EVs was used for sorting following the gating strategy described in Figure 8. Both populations of EV could be detected in the mixed sample, revealing positive events in gates P1, P2, P3, and P4 (Fig. 9A), confirming effective CFSE and antibody labeling of both EV populations and no overlap of signal.

**Figure 9.**
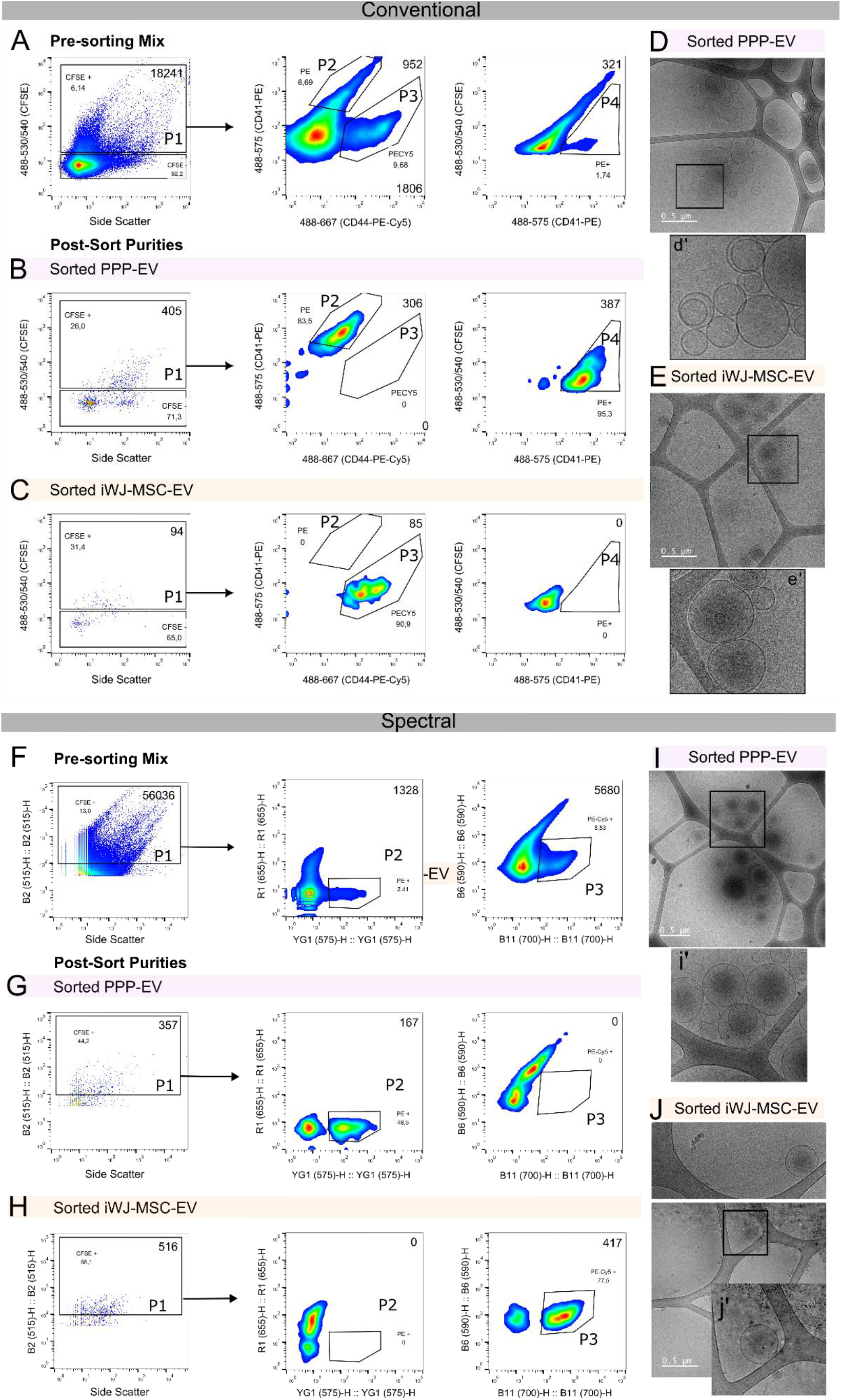
Sorting and characterization of antibody-labeled EVs by conventional and spectral Flow cytometry. (A) Flow cytometry plots of the mixed EV sample used for sorting, containing CD41-PE⁺ platelet-derived EVs (PPP-EVs) and CD44-PE-Cy5⁺ iWJ-MSC-derived EVs. (B-C) Post-sort purity analysis of the CD41-PE⁺ PPP-EV fraction (B) and CD44-PE-Cy5⁺ MSC-EV fraction (C). Event counts for 60 s of acquisition time are indicated within the plots: gates P1, P2, and P4 are shown in the top right corner while gate P3 counts are shown in the bottom right corner. (D-E) Cryo-transmission electron microscopy (Cryo-TEM) images of sorted PPP-EV (D) and iWJ-MSC-EV (E) populations confirmed the presence of intact, nanosized vesicles. (F) Spectral flow cytometry plots of the mixed EV sample used for sorting, containing CD41-PE⁺ platelet-derived EVs (PPP-EVs) and CD44-PE-Cy5⁺ iWJ-MSC-derived EVs. (G-H) Post-sort purity analysis of the CD41-PE⁺ (PPP-EVs) (G) and CD44-PE-Cy5⁺ (MSC-EVs) sorted populations (H). (I-J) Cryo-TEM images of the PPP-EV (I) and MSC-EV (J) sorted populations confirming the presence of intact, nanosized vesicles.

After sorting P2 (CD41-PE⁺) and P3 (CD44-PE-Cy5⁺) populations, we obtained two highly pure populations corresponding to CD41⁺ (PPP-EVs) and CD44⁺ events (iWJ-MSC-EVs) (Fig. 9B and C). The post-sort CD41⁺ PPP-EV sample showed events exclusively in the P2 and P4 control gates (95.3%), validating the specificity of the PE signal and confirming correct identification and isolation of CD41⁺ EVs (Fig. 9B). Also, the post-sort CD44⁺ MSC-EV sample displayed positive events only in the P3 gate (90.9%), consistent with specific CD44 labeling and sorting (Fig. 9C). The presence of nanosized, round-shaped, double-membraned vesicles in both sorted populations was confirmed by cryogenic transmission electron microscopy (Cryo-TEM) (Fig. 9D and E), supporting the successful isolation of intact EV subtypes.

To determine whether distinct EV subtypes labeled with specific antibodies could be successfully sorted in the spectral system, the same mixed sample of CFSE-labeled CD41-PE⁺ PPP-EVs and CD44-PE-Cy5⁺ MSC-EVs was used for sorting following the gating strategy described in Fig 8. The starting mixed sample revealed positive events in gates P1, P2, P3, and confirmed effective CFSE staining and antibody labeling of both EV populations (Fig. 9F). Sorting resulted in two highly pure populations corresponding to CD41⁺ PPP-EVs and CD44⁺ MSC-EVs (Fig. 9G, H). The post-sort PPP-EV fraction contained events exclusively within the P2 gate, clearly differentiating from background, and confirming accurate identification of CD41⁺ EVs (Fig. 9G). Conversely, the post-sort MSC-EV fraction showed positivity only in the P3 gate, consistent with specific CD44 labeling (Fig. 9H). The presence of nanosized, round-shaped, double-membraned vesicles in both sorted fractions was confirmed by cryogenic transmission electron microscopy (Cryo-TEM) (Fig. 9I and J), supporting the successful isolation and structural integrity of the sorted EV subtypes.

As one of the distinctive capabilities of the spectral cell sorter is its integrated CellView imaging technology, we further examined representative images and fluorescence spectra of the sorted EV populations. This feature enabled direct visual confirmation of detected particles in real time, providing additional evidence that the sorted events corresponded to the desired fluorescent phenotype rather than coincident noise or aggregates (Suppl. Fig. 3). The CellView images of both PPP-EVs and MSC-EVs displayed the expected fluorescence profiles consistent with their respective CFSE and antibody labels. This imaging-based verification complements conventional scatter and fluorescence data, enhancing confidence in the specificity and purity of the sorted EV fractions and offering a valuable layer of quality control in single-vesicle sorting workflows.

### EV sorting using both conventional or spectral cytometers yielded sufficient quantities of EV RNA to perform small RNA sequencing analyses

One of the main objectives of applying flow cytometry and EV sorting in the EV field is to enable the molecular characterization of specific EV subpopulations. In this context, we performed small RNA extractions from the sorted populations either from the conventional or spectral sorters. The extracted RNA was then quantified to assess whether the yield and quality was sufficient for downstream small RNA sequencing analysis (RNAseq) (Fig. 10A).

**Figure 10.**
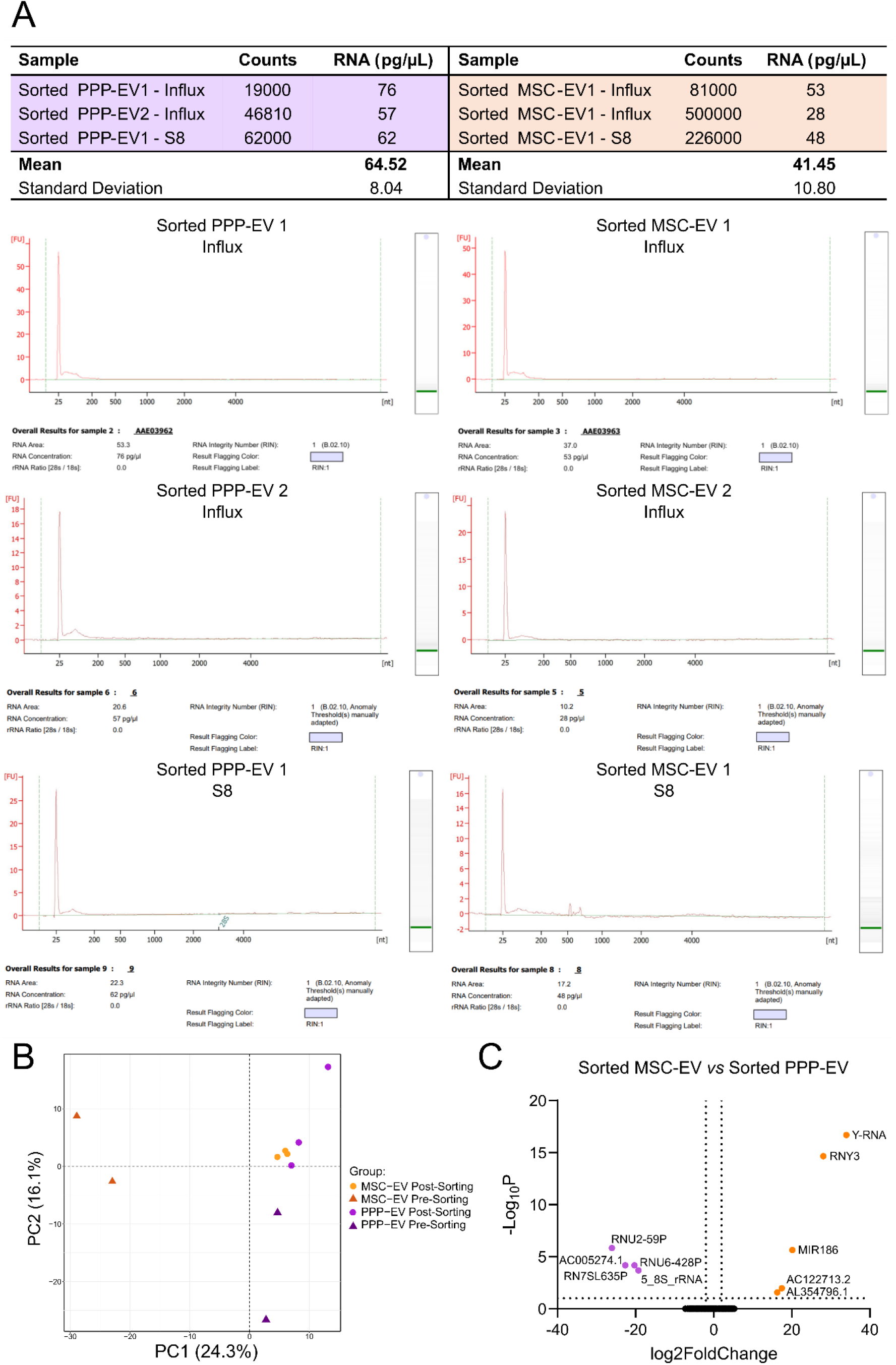
RNA quantification and small RNA sequencing analysis of sorted PPP-EVs and MSC-EVs. (A) Small RNAs were extracted from the sorted populations of CD41⁺ PPP-EVs and CD44⁺ iWJ-MSC-derived EVs obtained from three independent sorting experiments. The table presents the EV counts obtained from each sorting and the amount of extracted RNA. Representative Bioanalyzer electropherograms from the six post-sorting samples are presented in the bottom, showing RNA yields and quality check. (B) Principal component analysis (PCA) of small RNA profiles from pre- and post-sort fractions of PPP-EVs and iWJ-MSC-derived EVs, showing clustering based on sample origin and sorting status. (C) Volcano plot depicting differentially expressed small RNAs between sorted PPP-EV and MSC-EV populations. Significantly upregulated and downregulated small RNAs are highlighted, illustrating the distinct molecular signatures of each EV population.

RNA quantification analysis demonstrated sufficient yield in all post-sort fractions for downstream RNA sequencing, with concentrations ranging from 28 to 76 pg/μL (Fig. 10A). Notably, as few as ∼ 2x10^5^ sorted EVs provided enough RNA for sequencing. These amounts were obtained within relatively short sorting times, ranging from 3-5 h for the conventional sorter to 1-2 h for the spectral sorter. However, RNA amounts varied between sorting experiments independently from the cytometer used, with standard deviations of 8.04 pg/μL for PPP-EVs and 10.80 pg/μL for MSC-EVs. Notably, RNA yield did not correlate with the number of particle counts recorded during sorting, suggesting that factors beyond event counts, such as EV integrity, RNA cargo variability, or sorter-specific recovery efficiency may influence RNA recovery.

To confirm that the RNA content obtained from the post-sort fractions was suitable for molecular analysis, we conducted a pilot small RNA sequencing study (Fig. 10B and C). The results demonstrated the feasibility of downstream transcriptomic analysis and enabled the identification of specific small RNAs in the sorted CD41^+^ PPP-EV and CD44^+^ MSC-EV populations. Principal component analysis (PCA) clearly clustered pre-sorted MSC-EVs and PPP-EVs separately, indicating very distinct RNA profiles. However, sorted MSC-EV and PPP-EV populations exhibited distinguishable, yet much closer profiles, suggesting a few differentially expressed small RNAs between them (Fig. 10B). Despite this, differential expression analysis revealed specific small RNAs that remained differentially expressed between the post-sort PPP-EV and MSC-EV samples (Fig. 10C), supporting the retention of molecular identity following sorting.

## Discussion

In this study, we established and optimized a fluorescence-based nanoFACS workflow for the detection, phenotyping and physical isolation of defined extracellular vesicle (EV) subpopulations. Importantly, the workflow was successfully implemented on two distinct sorting platforms, a conventional BD Influx™ and a spectral BD FACSDiscover™ S8, and generated highly enriched EV populations while preserving vesicular integrity and compatibility with downstream small RNA sequencing. These findings support nanoFACS as a strategy not only for analyzing EV heterogeneity but also for physically isolating phenotypically defined populations for subsequent molecular characterization.

One of the main challenges in EV flow cytometry remains the reliable detection of nanoscale structures, whose small size and low refractive index place their optical signals close to the detection limits of conventional cytometers (26,27). At this scale, electronic noise, free dyes, protein aggregates and buffer contaminants can generate signals overlapping with genuine EV-associated events. Instrument calibration and appropriate reference materials are therefore particularly important. In our study, the combined use of non-fluorescent NIST particles, fluorescent reference particles and Rosetta beads provided complementary information on scatter and fluorescence performance. This strategy is consistent with current recommendations for small-particle flow cytometry, where standardization of instrument sensitivity and fluorescence-based detection can improve discrimination from background (19). Importantly, equivalent diameters obtained using FCM_PASS_ should not be interpreted as direct measurements of physical EV size. Because EVs have substantially different refractive indices and optical properties from polystyrene calibration beads, these values represent standardized optical-equivalent diameters rather than actual vesicle diameters.

The triggering strategy is closely related to these detection limitations. Although optimized FSC- or SSC-based approaches have successfully detected nanoscale particles under specific instrumental conditions (28–30), scatter sensitivity decreases substantially for small particles with low refractive index. Some studies have demonstrated that unstained EV-enriched preparations can be distinguished from electronic background using highly optimized instrument settings using the violet laser (9). However, fluorescence labelling is still commonly required when EV populations need to be specifically identified and subsequently sorted, in agreement with our observations and previous studies (31). Thus, rather than relying exclusively on scatter, our results reinforce the importance of robust fluorescence labelling, stringent controls and appropriate acquisition conditions for reliable EV sorting, consistent with previous reports (32,33), MiFlowCyt-EV standards (13) and MISEV recommendations (17,18).

In this context, CFSE provided a useful fluorescence-based strategy for detecting membrane-enclosed particles. Given the intrinsically low fluorescence expected from EVs, sufficiently bright fluorophores detectable through readily accessible optical channels were prioritized to maximize signal recovery across instruments. CFSE has previously been used successfully for nanoFACS because it enters membrane-enclosed structures and reacts with intraluminal amines, favoring retention of the fluorescent signal (34). In our study, CFSE-labelled EV-enriched SEC fractions were clearly separated from unstained EVs and from control samples, including protein-rich SEC fractions and CFSE-treated cell-free medium, on both platforms. Furthermore, RIPA-mediated membrane disruption markedly reduced the CFSE-positive signal, supporting its association with membrane-enclosed particles. Nevertheless, CFSE positivity alone cannot establish EV identity. Rather, these controls, together with EV enrichment and subsequent characterization, such as cryo-TEM imaging, provide converging evidence that the detected populations were enriched in vesicular structures.

Fluorescence labelling of surface markers subsequently enabled phenotypically distinct EV subsets to be independently sorted. Previous nanoFACS studies have demonstrated that fluorescence-based labelling can resolve specific EV populations (9,31,34), and our findings further show that these populations can be recovered with high purity on both conventional and spectral sorting platforms. An additional practical aspect of our workflow was the removal of free antibody by rapid ultrafiltration. Compared with approaches in which an additional SEC purification step is performed after fluorescent labelling (34), ultrafiltration avoided an additional chromatographic separation, thereby reducing processing time and potential sample loss while maintaining sufficient EV recovery for sorting. This is particularly relevant in EV applications, where starting material and final yields are often limited.

Beyond staining, sample concentration and acquisition conditions strongly influence nanoFACS performance. Flow rate and sample dilution were critical in our experiments, consistent with previous studies showing that high particle concentrations increase coincident detection or “swarming” (10). Under these conditions, multiple particles may simultaneously cross the interrogation point and be recorded as a single event, potentially altering apparent event counts and fluorescence intensity. Conversely, excessive dilution reduces event recovery and substantially increases sorting time. Our findings therefore support empirical optimization of dilution and flow rate rather than assuming that maximal dilution or the lowest possible flow rate will necessarily provide optimal conditions.

The behavior of the protein-rich control fractions further illustrates the influence of sample composition on nanoscale flow cytometry. These fractions required greater dilution than EV-enriched samples before their signal could be adequately distinguished from background. This observation is consistent with the dilution paradox described by Rops et al. (35) in heterogeneous samples such as plasma and cell culture supernatants, where changes in sample concentration can disproportionately affect background and coincident events. Based on these observations, we used consistent dilution conditions for samples originating from the same biological matrix and recommend appropriate EV enrichment or purification before nanoFACS analysis of highly complex specimens.

Although the general methodological requirements were shared between platforms, the comparison of the conventional and spectral sorters revealed relevant differences. Both instruments successfully isolated the targeted EV populations with high post-sort purity and preserved vesicle integrity, demonstrating that the proposed workflow is not restricted to a single instrument architecture. However, under our experimental conditions, the spectral sorter provided narrower fluorescence distributions and clearer separation between phenotypically defined populations, with no detectable non-specific PE signal in the corresponding controls. The improved resolution may partly reflect differences in optical and detector architecture, including the use of avalanche photodiodes (36), although our data do not allow the contribution of individual hardware components to be established.

Operational differences were also evident. The spectral sorter generated sufficient post-sort material for downstream molecular analysis within approximately 1–2 h, whereas the conventional platform required approximately 3–5 h to obtain comparable yields. Despite these differences, both instruments performed considerably faster than previously reported sorting approaches (9) and generated sufficient material for subsequent analysis. Because the two cytometers differ substantially in their optical and fluidic configurations, their acquisition settings cannot be considered identical. Rather, comparable experimental conditions and equivalent gating principles were applied. Overall, the BD Influx™ offered extensive flexibility in acquisition configuration, whereas the FACSDiscover™ S8 provided greater fluorescence resolution and faster recovery under the conditions tested. The discontinuation of the BD Influx™ additionally limits its future implementation and highlights the importance of transferring validated nanoFACS principles to newer generation instruments.

A major strength of the workflow is that validation extended beyond cytometric purity. Fluorescence-positive events alone cannot unequivocally establish that the recovered material corresponds to structurally intact and biologically meaningful EVs. Cryo-TEM therefore provided an important orthogonal validation by demonstrating preservation of vesicular structures after sorting. This structural evidence was further complemented by downstream molecular analysis, strengthening the interpretation of the sorted fractions beyond fluorescence detection alone.

Despite the relatively low amount of recovered material, RNA concentrations ranging from 28 to 76 pg/µL were sufficient for library preparation and small RNA sequencing. This is consistent with previous studies demonstrating the feasibility of transcriptomic analysis from low amounts of EV-derived RNA (37,38). Interestingly, RNA yield did not correlate directly with cytometric event counts. This may reflect biological heterogeneity in RNA cargo between EV populations, variation in extraction efficiency or technical variability associated with low-input RNA, reinforcing previous recommendations for stringent pre-analytical and analytical standardization in EV-RNA studies (39).

Pilot small RNA sequencing provided an additional level of biological validation. While pre-sort PPP-EV and MSC-EV preparations showed clear separation by principal component analysis, post-sort fractions clustered more closely. Nevertheless, subtype-associated small RNAs remained detectable following sorting, supporting previous observations that distinct EV populations can retain molecular signatures associated with their source (39) and demonstrating the feasibility of combining nanoFACS with downstream molecular profiling. The reduced separation observed after sorting may reflect the low RNA input, loss of low-abundance transcripts during processing, enrichment of RNA species shared between the selected EV populations or greater molecular similarity between the phenotypically selected subsets than between the original bulk preparations. These sequencing results should therefore be interpreted primarily as a demonstration of downstream molecular feasibility rather than as a comprehensive characterization of subtype-specific transcriptomes.

Taken together, these results highlight the value of combining complementary validation strategies when applying nanoFACS to EV analysis. Neither CFSE staining, detergent sensitivity, antibody phenotyping nor high post-sort purity alone is sufficient to establish EV identity. Instead, EV enrichment, appropriate fluorescence controls, detergent sensitivity, surface-marker phenotyping, cryo-TEM and downstream molecular analysis provide multiple independent layers of evidence supporting the isolation of biologically meaningful EV-enriched populations.

Several limitations should nevertheless be acknowledged. First, the workflow was optimized using defined EV preparations and selected experimental matrices, and its performance in highly complex clinical samples will require further validation. Second, the small RNA sequencing analysis was exploratory and based on low RNA input and a limited number of sorted populations, while phenotypic isolation in the present study relied on a restricted number of surface markers.

Future developments should therefore focus on multiparametric phenotyping using combinations of surface markers rather than single-marker definitions. This may be particularly relevant for spectral flow cytometry, where higher-dimensional fluorescence analysis could enable more specific discrimination and isolation of EV subsets. Such approaches could facilitate the identification of rare populations in complex biological or clinical specimens and, importantly, allow the relationship between EV surface phenotype and molecular cargo to be investigated directly.

In summary, we established a fluorescence-based nanoFACS workflow capable of isolating highly enriched EV subpopulations while preserving vesicular integrity and compatibility with downstream molecular analysis. Both conventional and spectral sorting platforms were effective, although they displayed distinct analytical and operational characteristics. Across platforms, reliable EV sorting depended on appropriate fluorescence labelling, stringent controls, calibration and optimization of sample dilution and flow conditions. The combination of cytometric, structural and molecular validation provides a practical framework for the isolation and subsequent molecular characterization of phenotypically defined EV populations and for investigating EV heterogeneity.

## Supplementary Figures

**Supplementary Figure 1.**
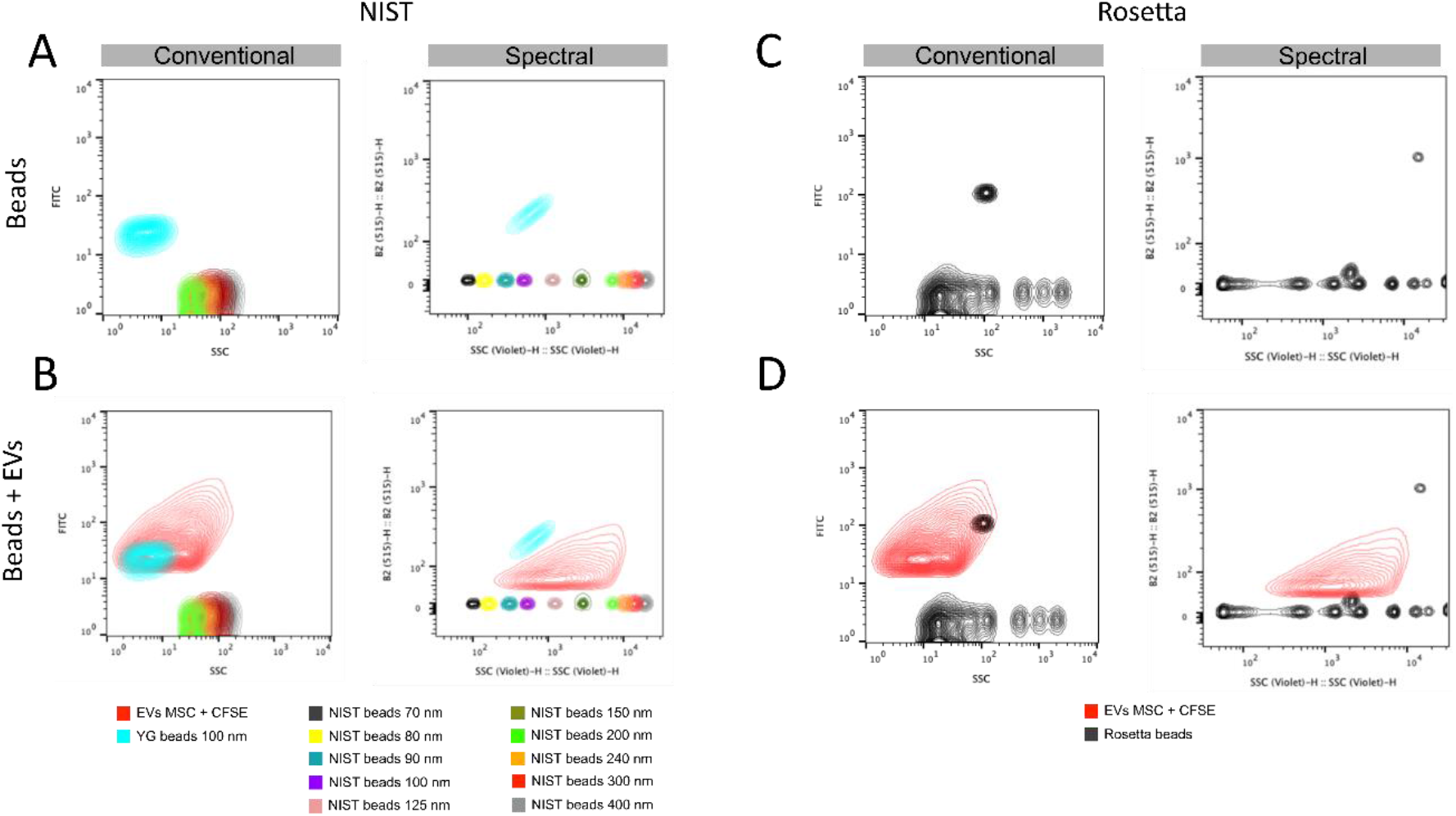
Fluorescence-triggered acquisition comparably identifies small EV in conventional and spectral cytometers. (A) SSC/FITC intensity distributions of NIST-traceable polystyrene beads together with 100-nm YG beads acquired on the conventional BD Influx (left) and spectral S8 cytometer (right). (B) SSC/FITC signal of fluorescence-triggered acquisition of CFSE-stained EVs overlaid on the NIST/YG beads signal, acquired either on the conventional (left) or spectral cytometer (right). Most EV events were distributed below an equivalent diameter of approximately 200 nm. (C) SSC/FITC signal of Rosetta calibration beads acquired on the conventional (left) and spectral cytometer (right). (D) SSC/FITC distribution of fluorescence-triggered acquisition of CFSE-stained EVs (red) overlaid to the Rosetta beads signal on the BD Influx (left, blue) and S8 (right, black).

**Supplementary Figure 2.**
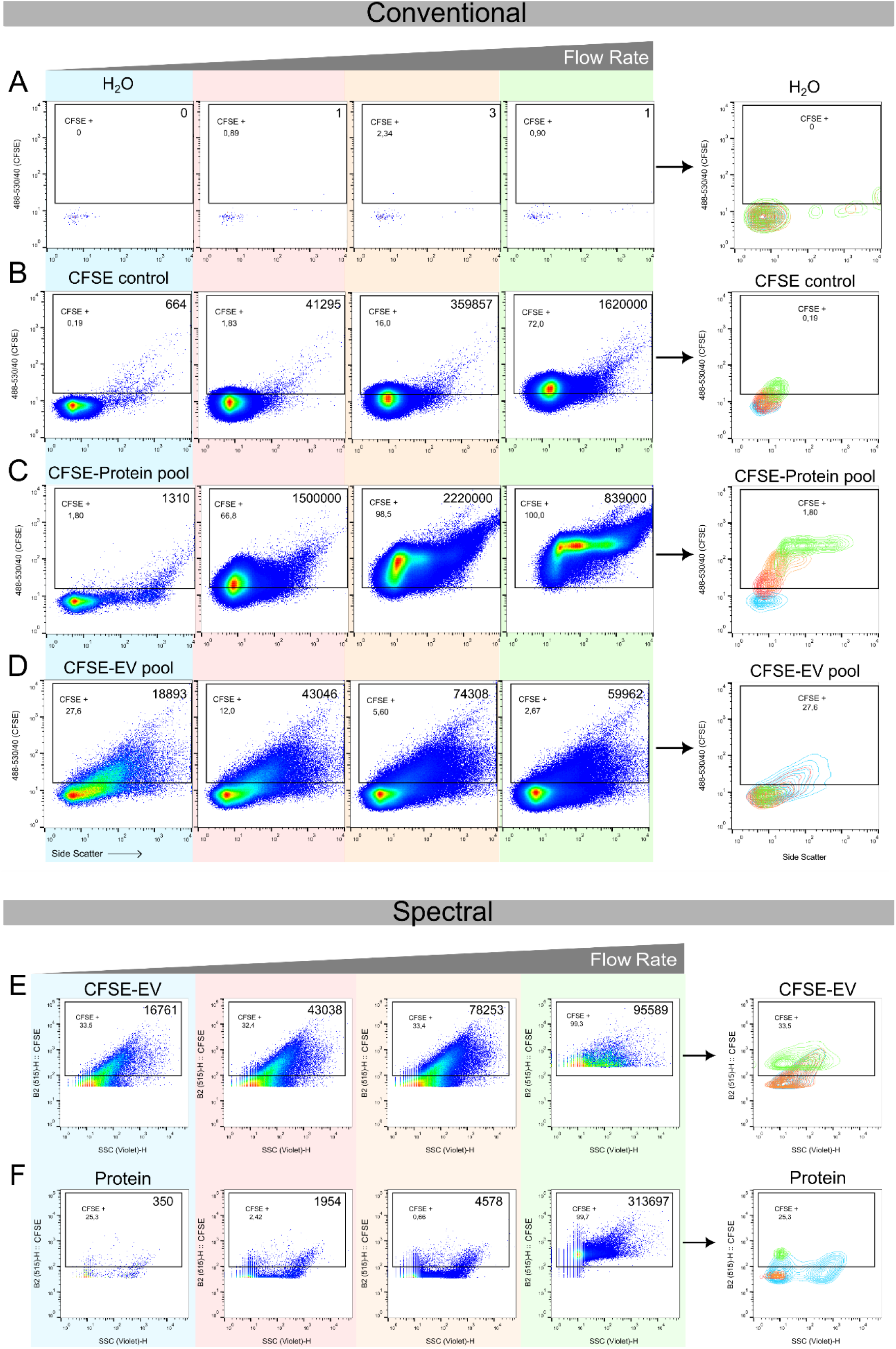
Flow rate increases signal artefacts and impacts resolution of EV detection by flow cytometry. BD Influx. (A-D). Representative flow cytometry plots for (A) 0.22 µm–filtered water, (B) CFSE-labeled cell-free medium control, (C) CFSE-labeled protein-enriched SEC pooled fraction, and (D) CFSE-labeled iWJ-MSC–EV pool. Samples were analyzed for 60 s at increasing flow rates corresponding to sheath pressures of 21.4, 21.5, 21.8, and 25 psi, depicted in blue, red, orange, and green, respectively. The CFSE⁺ count (60 sec) is indicated in the top right corner of each plot.**BD S8.** (E-F) Effect of flow rate on signal intensity in CFSE-labeled EVs (E) and protein-enriched fractions (F). Flow rates are represented in blue, red, orange, and green at 1, 20 and 50 relative flow rates, corresponding to 0.16, 0.31 and 0.75 µL/s respectively, illustrating the influence of flow rate on detection sensitivity and resolution. The final plot of each row is a superposition contour plot of all the flow rates represented. Event counts for each 60 s acquisition are shown in the top right corner of each plot.

**Supplementary Figure 3.**
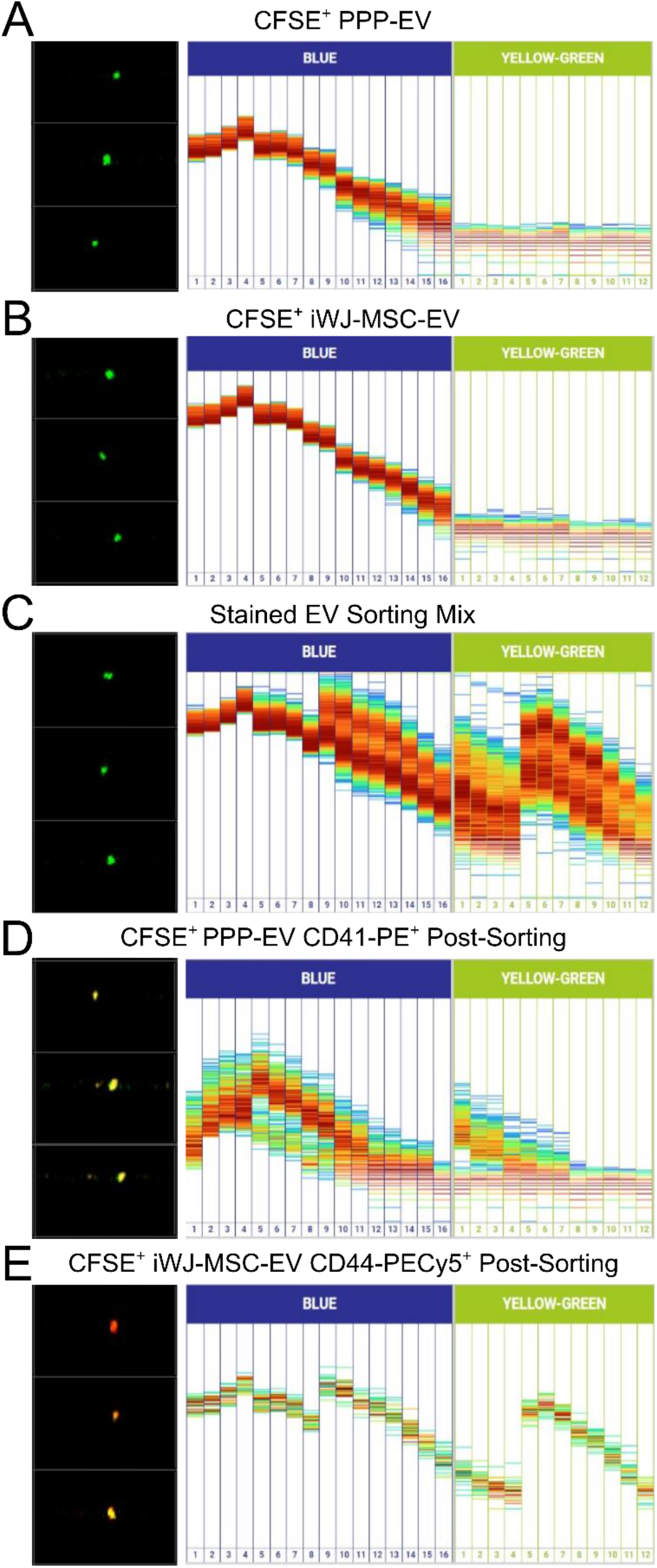
Representative CellView images and fluorescence spectra (blue and yellow-green channels) acquired using the BD FACSdiscover^TM^ S8 cell sorter. (A) Poor-platelet plasma–derived EVs (PPP-EVs) stained with CFSE. (B) Mesenchymal stromal cell–derived EVs (MSC-EVs) stained with CFSE. (C) Sorting mix containing both CFSE-stained PPP-EVs and MSC-EVs labeled with CD41-PE and CD44-PECy5 antibodies, respectively. (D) Post-sort PPP-EVs. (E) Post-sort MSC-EVs.

## Author’s contribution

**M.C.S.** and **M.M.T.**: Conceptualization, Methodology, Formal analysis, Data curation, Writing – original draft, Writing – review and editing; **S.G.G.**: Formal analysis, Data curation, Writing – review and editing; **E.J.**, **M.F.M.**, **E.R.**, **A.B.**, **N.B.**, **M.S.M.**, and **L.P.**: Methodology, Writing – review and editing; **B.F.** and **J.P.**: Formal analysis, Data curation, Writing – review and editing; **F.E.B.**: Conceptualization, Writing – review and editing; **O.F.** and **M.F.**: Conceptualization, Funding acquisition, Methodology, Formal analysis, Data curation, Writing – original draft, Writing – review and editing.

## Data availability

Flow cytometry is fully open and available in Figshare:

Fig2-3-4: https://doi.org/10.6084/m9.figshare.34054242

Fig5-9: 10.6084/m9.figshare.34056072

Fig10: 10.6084/m9.figshare.34054941

## Acknowledgements

We want to thank Martí de Cabo from Electron microscopy-Microscopy Service UAB.

